# Sticky Interactions Govern Sequence-Dependent Dynamics in Biomolecular Condensates

**DOI:** 10.1101/2025.07.09.664001

**Authors:** Keerthivasan Muthukumar, Dinesh Sundaravadivelu Devarajan, Young C. Kim, Jeetain Mittal

## Abstract

Intrinsically disordered proteins (IDPs) play a central role in shaping the dynamics and material properties of biomolecular condensates. Understanding how sequence features determine these properties is critical for elucidating physiological function and guiding the rational design of synthetic condensates. Here, we use molecular dynamics simulations to investigate condensates formed by model IDPs with systematically varied chain length and charge patterning, two features characteristic of natural IDPs. Our results show that chain relaxation times, governed by sequence-dependent electrostatic interactions, quantitatively predict condensate viscosity and diffusivity. These condensates exhibit dynamics consistent with a crossover regime between Rouse and reptation behavior. While the Rouse model with idealized friction fails to capture sequence effects, the sticky Rouse model, which incorporates transient interchain contact lifetimes, accurately predicts chain reconfiguration times and, consequently, macroscopic material properties. This work establishes a predictive, sequence-resolved framework that links molecular interactions to condensate dynamics across length and time scales.

## INTRODUCTION

Intrinsically disordered proteins (IDPs) undergo phase separation^1–4^ to form highly dynamic, liquid-like membraneless condensates that are central for cellular organization and biochemical processes^5,6^. The material properties of these condensates, such as viscosity^7,8^, and viscoelasticity^9–12^, underpin their stability and function, while the changes in these properties over time can signal the onset of neurodegenerative conditions such as amyotrophic lateral sclerosis and frontotemporal dementia^13,14^. These observations highlight the need to understand the sequence-encoded molecular determinants of condensate material properties. At the molecular level, the sequence characteristics of IDPs influence their interaction dynamics, directly shaping the material properties of condensates^15–19^. For example, arginine-rich IDPs form condensates with significantly higher viscosity than lysine-rich counterparts, due to stronger interaction networks^20,21^. Additionally, the patterning of charged residues in IDPs profoundly influences condensate properties. For instance, a diblock arrangement of charged residues promotes strong electrostatic attractions between oppositely charged segments, slowing dynamics^22^. Alterations in IDP sequences, whether by adjusting net charge or mutating glycine, aromatic, or arginine residues, reveal that diverse molecular interactions can govern condensate material properties^23^.

These observations motivate further investigation into how sequence-encoded features of IDPs modulate molecular interactions that shape condensate material properties. Notably, even in condensates with increased macroscopic viscosity, molecular-scale dynamics remain remarkably rapid^24^. This contrast raises key questions about which molecular interactions underlie macroscopic slowdowns in condensates. Previous research shows that transient residue-residue interactions, and the lifetimes of these contacts, are key determinants of condensate material properties^22^. Additional studies suggest that material properties are directly influenced by protein chain dynamics by accounting for the role of polymer entanglements^21^.

For multivalent associative proteins with complementary interactions, bond relaxation slows noticeably when stoichiometric balance limits alternative binding options^25^. A similar effect is seen in weakly interacting condensates, where dynamics are governed not by bond lifetimes but by the availability of binding stickers^26^. Studies of protein-nucleic acid condensates show that intermolecular cross-links formed by sequence-specific DNA hybridization can alter diffusivity and viscosity by modulating chain dynamics^27^. These findings collectively highlight that underlying molecular interactions dictate the macroscopic material properties of condensates. Although polymer theories suggest that chain length strongly affects material properties across scales^28,29^, the combined influence of IDP chain length and sequence features on condensate properties remains poorly understood. Given the inherent variability in natural IDP chain lengths^30^, it is important to understand how these differences affect sequence-dependent condensate behavior.

To address these questions, we investigate how chain length and charge patterning in IDPs influence dynamics across multiple length scales, with the goal of uncovering the molecular origins of condensate material properties such as chain diffusivity and viscosity. We also evaluate the applicability of polymer theory frameworks for quantitatively predicting the material properties of IDP condensates. To this end, we use the coarse-grained hydropathy scale (HPS) model to simulate charge-neutral model IDP condensates composed of equal numbers of glutamic acid (E) and lysine (K) residues. These model IDPs allowed us to systematically vary charge patterning, from perfectly alternating to diblock arrangements, covering a broad spectrum of charge segregation. By also varying chain length, we mimicked key features of natural IDPs, enabling us to study dynamics across multiple length scales and relate them to condensate material properties.

## RESULTS

The IDP condensates can be described through a multiscale framework that connects sequence features, microscopic dynamics, and macroscopic material properties (**Figure 1**). We investigated how two key sequence parameters, chain length 𝑁 and degree of charge segregation influence the dynamics of individual chains across multiple length scales, which in turn influence the macroscopic material properties of the dense phase. To enable comparison across different IDP lengths and charge patterning, we normalized the previously developed sequence charge decoration SCD parameter^31,32^ (nSCD) to range between 0 and 1 (with the lower and upper bounds corresponding to perfectly alternating and diblock sequences, respectively). By combining analyses at the monomer, chain, and condensate level, we sought to build a predictive understanding of how molecular-level features shape the macroscopic dynamics and rheology of IDP condensates.

**Figure 1:**
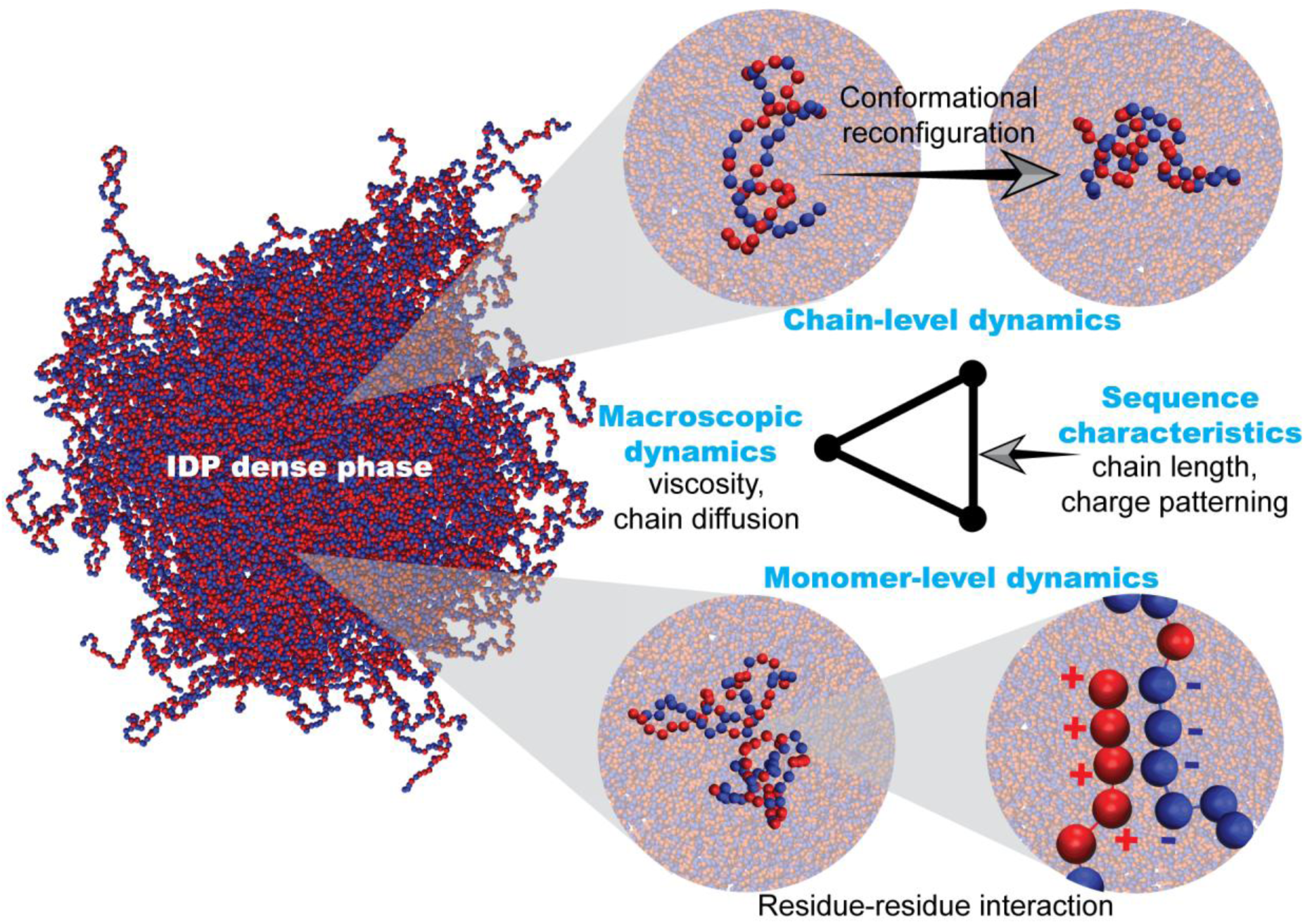
Multiscale analysis for understanding condensate material properties: Schematic illustrating the interplay between sequence features and dynamics at different length scales within the dense phase of condensates. Key sequence features such as chain length and charge patterning influence monomer-level to chain-level dynamics, which in turn dictate the macroscopic material properties of IDP condensates, highlighting the interdependence across different length scales.

### IDPs Adopt Ideal-Chain Conformations in the Dense Phase Regardless of Charge Patterning

The conformations of IDP chains play an important role in determining condensate dynamics and function^33–35^. We quantified chain conformations using the mean radius of gyration 𝑅_g_, computed from the gyration tensor 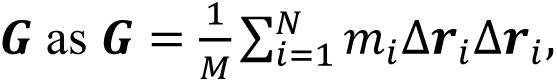 where 𝑚*_i_* is the mass of the 𝑖-th residue, Δ𝒓_𝑖_ is the displacement vector from the chain’s center of mass, and 𝑀 is the total mass of the chain. The radius of gyration was then obtained as 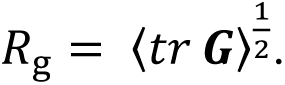. We observed that 𝑅_g_ increased with chain length according to the scaling law 𝑅_g_ ∼ 𝑁^𝜈^, with 𝜈 ≈ 0.5 across all nSCD values (**Figure 2**). This scaling exponent, characteristic of ideal chain behavior^28,29^, indicates that E-K sequences in the dense phase adopt similar conformations. Consistent with this, the mean end-to-end distance 𝑅_e_ (**Supplementary Figure S1**) followed the theoretical expectation for ideal chains: 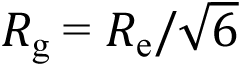 (**Figure 2 inset**). Similarly, the average interresidue distance 𝑅*_ij_* (**Supplementary Figure S2**) scaled with residue separation as |𝑖 − 𝑗|^0.5^, further supporting ideal chain-like behavior in the dense phase. Importantly, chain conformations showed minimal dependence on nSCD, consistent with previous findings at fixed chain length^33^. This insensitivity to sequence charge segregation contrasts with the dilute phase, where increasing nSCD promotes chain compaction due to stronger intrachain electrostatic interactions^22,33,36–38^. These results demonstrate that in the dense phase, IDPs adopt conformations closely resembling ideal chains, with structural variation governed primarily by chain length rather than sequence patterning.

**Figure 2:**
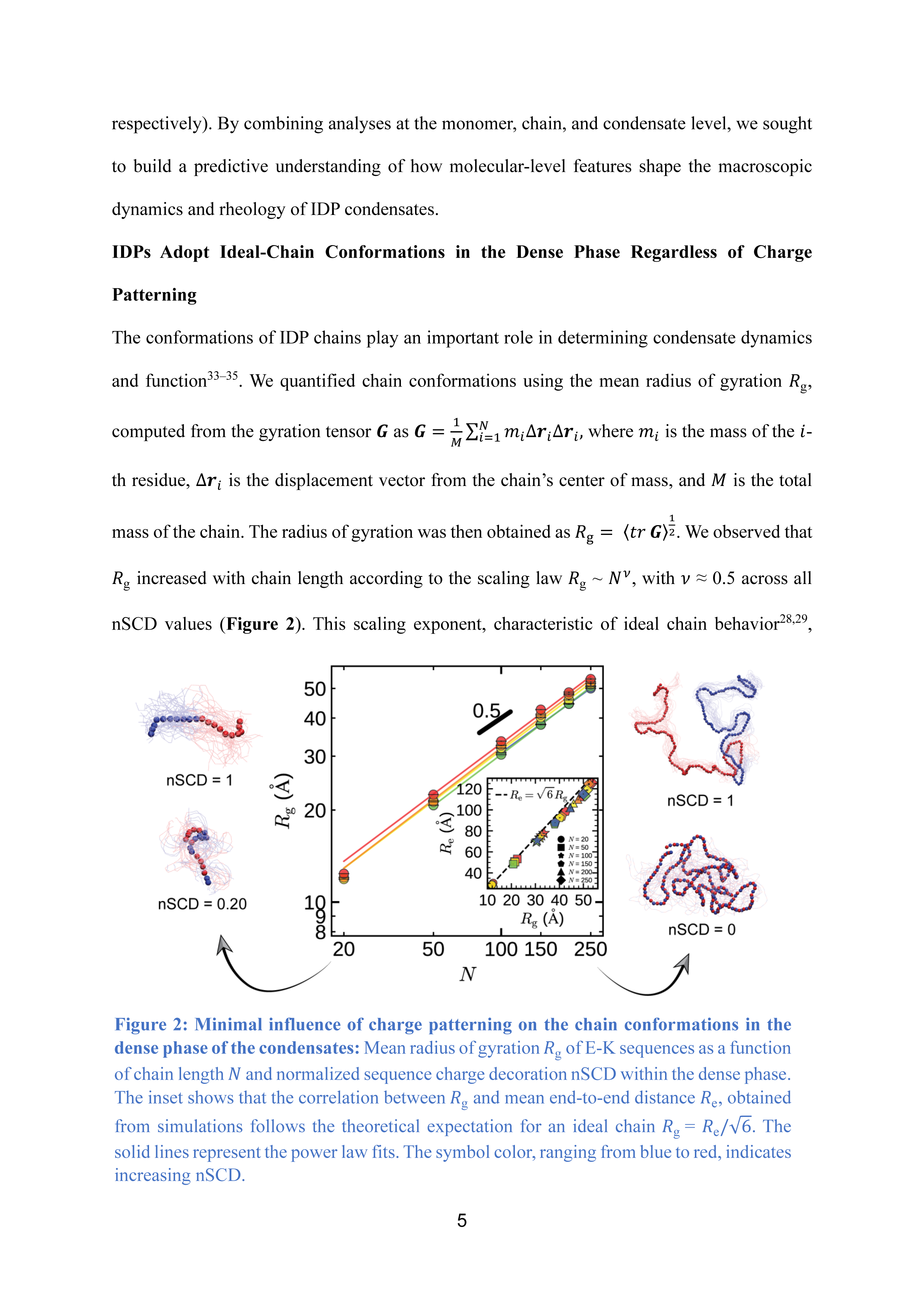
Minimal influence of charge patterning on the chain conformations in the dense phase of the condensates: Mean radius of gyration 𝑅_g_ of E-K sequences as a function of chain length 𝑁 and normalized sequence charge decoration nSCD within the dense phase. The inset shows that the correlation between 𝑅_g_ and mean end-to-end distance 𝑅_e_, obtained from simulations follows the theoretical expectation for an ideal chain 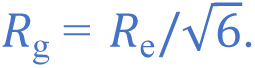 The solid lines represent the power law fits. The symbol color, ranging from blue to red, indicates increasing nSCD.

### Chain Length and Charge Patterning Modulate the Dynamics and Rheology of IDP Condensates

Having established how sequence features influence chain conformations in the dense phase, we next examined their impact on material properties. We focused on translational diffusivity

𝐷, a key measure of molecular mobility within the dense phase. To quantify chain motion, we computed the mean square displacement (MSD) of residues across dense phases formed by E-K sequences with varying 𝑁 and nSCD (**Supplementary Figure S3**). MSD was defined as MSD(𝑡) = 〈[𝒓_𝑖_(𝑡) − 𝒓_𝑖_(0)]^2^〉, where 𝒓_𝑖_(𝑡) is the position of residue 𝑖 at time 𝑡. Initially, we found that all sequences exhibited ballistic motion, with MSD scaling as 𝑡^2^. As interchain interactions became more pronounced, MSD transitioned to a subdiffusive regime with MSD ∼ 𝑡^0.5^. At longer times, the motion became diffusive with MSD growing linearly (MSD ∝ 𝑡^1^). These transitions occurred more slowly in sequences with longer 𝑁 and higher nSCD, indicating that these sequence features modulate chain dynamics. Polymer theories predict a decrease in diffusivity 𝐷 with increasing chain length 𝑁. In the Rouse model for unentangled polymers, 𝐷 ∼ 𝑁^−1^. In contrast, entangled polymer systems exhibit 𝐷 ∼ 𝑁^−2.3^ in experiments, approximately consistent with the theoretical expectation for reptation dynamics (𝐷 ∼ 𝑁^−2^)^28,29^. To test this scaling behavior in the E-K condensates, we extracted 𝐷 values from the diffusive regime using the relation MSD = 6𝐷𝑡 (**Supplementary Figure S4**). For a given 𝑁, increasing nSCD monotonically reduced 𝐷, due to stronger interchain electrostatic interactions arising from charge segregation^22,38^. Across different nSCD values, 𝐷 decreased with increasing 𝑁, following a power law with exponents ranging from-1.70 to-2.24 (**Figure 3a**). These exponents suggest that the system is mostly in a crossover regime between Rouse and reptation behavior. Additionally, only at high nSCD and longer 𝑁, normalized MSD values (MSD/t^0.5^; **Supplementary Figure S5**) showed a gradual decrease in slope values from 0 (Rouse-like) toward-0.25 (reptation-like) in the intermediate regime, before reaching diffusive motion with a slope of 0.5^36^. This indicated that systems are close to the onset of reptation-like behavior only at higher nSCD and longer *N*^36^. From these observations, we concluded that both 𝑁 and nSCD modulate molecular diffusion in the IDP dense phases.

**Figure 3:**
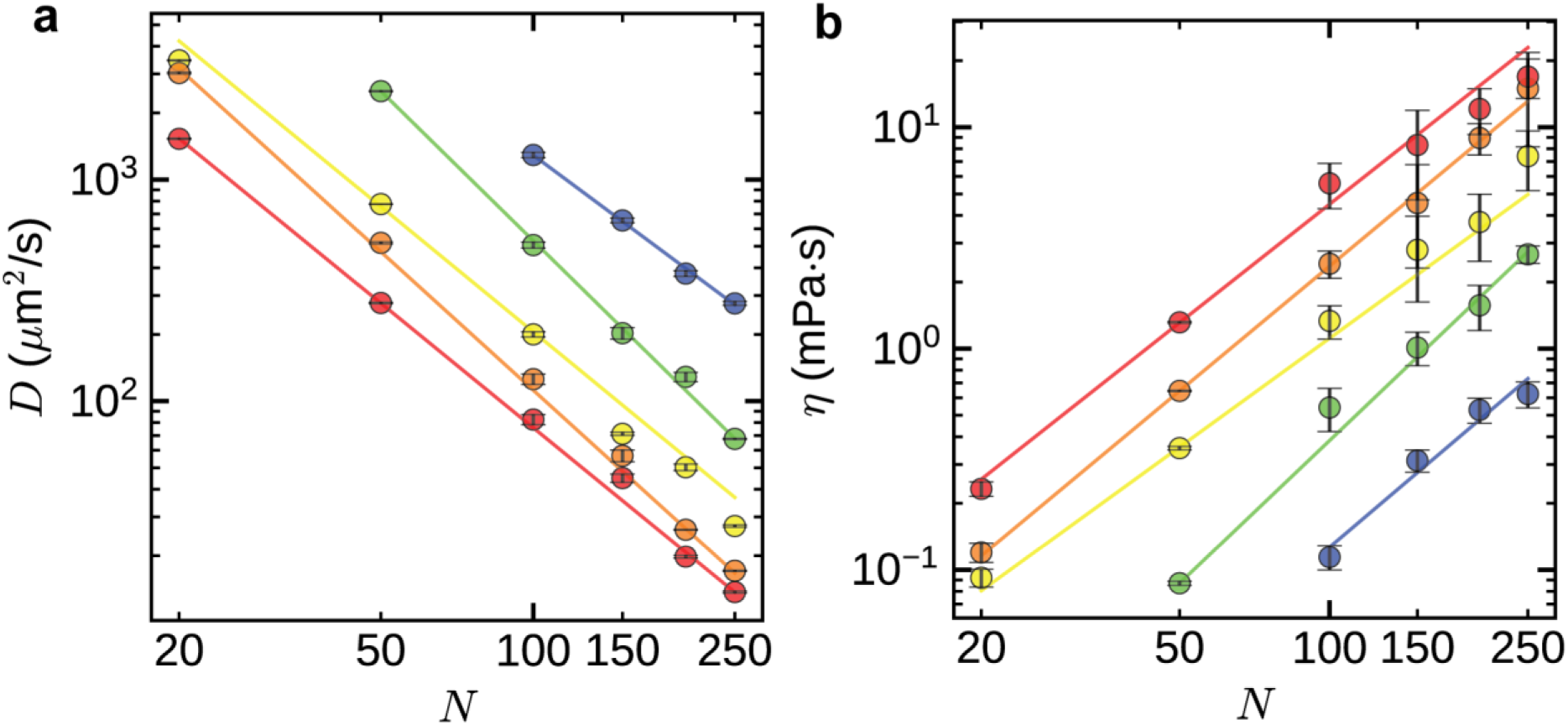
Longer and charge-segregated IDP chains form dynamically slower condensates: Material properties of the dense phase as a function of chain length 𝑁 and normalized sequence charge decoration nSCD: (a) Chain diffusivity 𝐷 and (b) Viscosity 𝜂. The solid lines represent the Bayesian power law fits. The symbol color, ranging from blue to red, indicates increasing nSCD.

We next computed the viscosity 𝜂 of dense phases formed by E-K sequences with varying 𝑁 and nSCD, using the Green-Kubo relation (see Methods). Polymer theories predict 𝜂 ∼ 𝑁^1^ in the Rouse model^28,29^, while experimental results in the reptation regime show 𝜂 ∼ 𝑁^3.4^. In our simulations, 𝜂 increased with 𝑁 following a power law, with exponents between 1.91 and 2.14 across different nSCD values (**Figure 3b**). These values are steeper than predicted by the Rouse model but remain well below those associated with fully reptating polymers. These results suggest that our simulated condensates lie in a crossover regime, where increasing chain length and interchain interactions begin to influence rheology without yet producing fully entangled dynamics. Although viscosity estimates at longer 𝑁 show larger uncertainties due to statistical noise inherent in the Green–Kubo method, which has been widely used to characterize condensate viscosities^39–41^ we believe that the scaling trends remain robust. For a given *N*, increasing nSCD also led to higher 𝜂 values, reflecting enhanced interchain electrostatic interactions (**Figure 3b**). Moreover, the data collapsed onto a single master curve following 𝐷 ∼ 𝜂^−1^ (**Supplementary Figure S6**), confirming the validity of the Stokes-Einstein relation in the dense phase. Together, these results highlight the critical role of sequence features such as chain length and charge patterning in shaping the material properties of the dense phase. The scaling exponents further support that the condensates are in a crossover regime between Rouse-like and reptation-dominated dynamics.

### Normal Mode Analysis Reveals Multiscale Relaxation in IDP Dense Phases

To investigate the molecular origins of condensate material properties, we performed a normal mode analysis. This approach characterizes relaxation dynamics across different segmental lengths, from monomers to entire chains^36,42–45^. We computed the autocorrelation function 𝑐_𝑝_ of the normal modes 𝑿_𝑝_ within the dense phase (see Methods), where higher mode number 𝑝 correspond to smaller chain segments (**Figure 4a**). We found that 𝑐_𝑝_ for different 𝑁 and nSCD decayed to zero (**Supplementary Figure S7**), indicating complete relaxation. We extracted relaxation times 𝜏_𝑝_ by fitting 𝑐_𝑝_ to a stretched exponential form (**Figure 4b and Supplementary Figure S8**). As expected, low-𝑝 modes, which represent chain-level dynamics, relaxed more slowly, whereas high-𝑝 modes corresponding to smaller segments relaxed more rapidly.

**Figure 4:**
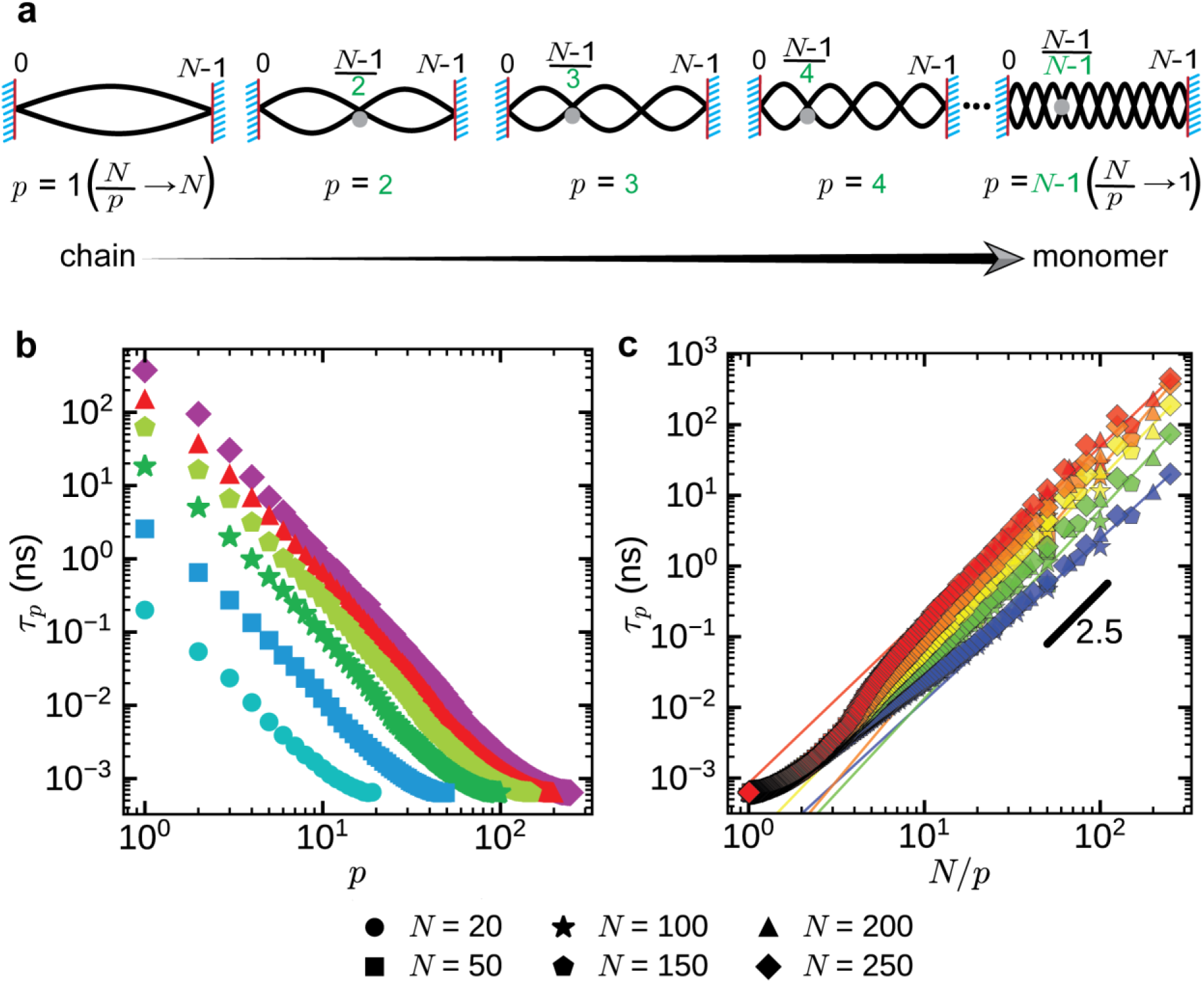
Normal modes reveal sequence-dependent multiscale dynamics in IDP condensates: (a) Schematic representation of normal modes of increasing mode number 𝑝 for a polymer chain of length 𝑁. Mode 𝑝 **=** 1 corresponds to whole-chain motion, while higher modes (*p* = 2, 3,…) represent increasingly localized segmental fluctuations down to the monomer scale (𝑝 = 𝑁–1). (b) Normal mode relaxation time 𝜏_𝑝_ of E-K sequences with normalized sequence charge decoration nSCD = 0.46 as a function of mode number 𝑝 and chain length 𝑁 within the dense phase. (c) 𝜏_𝑝_ as a function of normalized mode number 𝑁/𝑝 for different 𝑁 and nSCD within the dense phase. The solid lines in (c) represent the power law fits. The symbol color in (b), ranging from blue to magenta, indicates increasing 𝑁 and those in (c), ranging from blue to red, indicates increasing nSCD.

Segments at the monomer scale (𝑝 ≈ 𝑁 − 1) relaxed quickly and showed no significant variation with 𝑁 (**Figure 4b and Supplementary Figure S8**). Segments spanning up to approximately three monomer units exhibited minimal changes in relaxation across different nSCD values (**Figure 4c and Supplementary Figure S9**). For longer segments, relaxation times 𝜏_𝑝_ increased with both 𝑁 and nSCD, mirroring the slowdown observed in diffusivity and viscosity. When plotted against normalized mode number 𝑁/𝑝, segmental relaxation times for different 𝑁 collapsed onto a single curve at each nSCD value. The data followed a power-law scaling τ_𝑝_ ∼ (𝑁/𝑝)^𝜈^, with 𝜈 ranging from approximately 2.2 to 2.9 (**Figure 4c**). Again, these exponents indicate that segmental dynamics fall within the crossover regime between Rouse (2) and reptation (3.4) scaling^28,29^.

Overall, the normal mode analysis revealed that increasing 𝑁 and nSCD slowed the relaxation of several intermediate-sized segments, mirroring the dynamic slowdown observed in the condensate material properties. Because our simulation results place the E-K condensates in the crossover region between Rouse and reptation regimes, we next examined whether polymer dynamics theories, starting with the Rouse model, could quantitatively describe the trends observed in condensate properties.

### Rouse Theory Fails to Capture Medium Friction in IDP Dense Phases

The Rouse model describes a polymer chain as a series of 𝑁 beads connected by harmonic springs with root mean square length 𝑏. Each bead undergoes Brownian motion and experiences a friction coefficient 𝜁 (**Figure 5a**)^21,28,29,46^. No other interactions between the beads are included. Chain reconfiguration dynamics can be characterized by the relaxation time 𝜏_R_ of the end-to-end vector 𝑹_e_, which for a Rouse chain is given by:

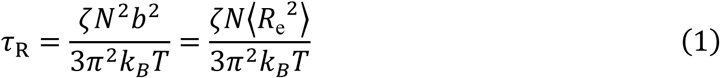

**Figure 5:**
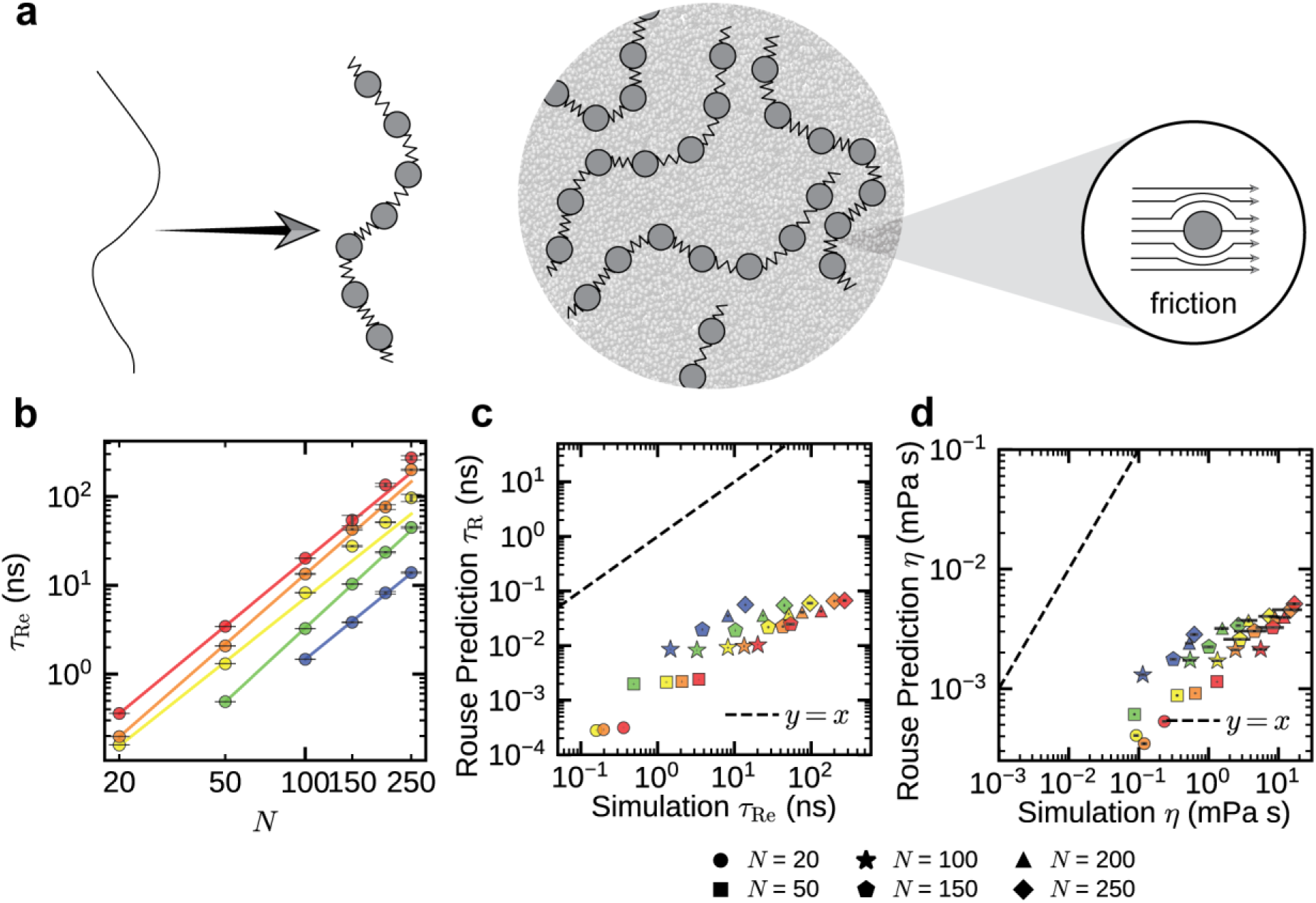
Rouse theory does not account for the medium friction experienced by the charged residues within the dense phase: (a) Schematic representation of the Rouse model where a polymer is modeled as beads connected by springs, exhibiting random motion and slowed by fluid friction. (b) End-to-end vector relaxation time 𝜏_Re_ computed from the simulated dense phase of E-K sequences as a function of chain length *N* and normalized sequence charge decoration nSCD. (c) Correlation plot comparing Rouse theory predictions and simulated end-to-end vector relaxation times where predictions were made using Equation (1), (d) Correlation plot comparing Rouse theory predictions and simulated dense phase viscosities where predictions were made using Equation (3). The solid lines in (b) represent the Bayesian power law fits, while dashed line in (c) and (d) corresponds to the 𝑦 = 𝑥 line. The symbol color, ranging from blue to red, indicates increasing nSCD.

where 𝑘_B_ is the Boltzmann constant, and 𝑇 is the temperature. The expression for 𝜏_R_ uses the ideal chain relation ⟨𝑅_e_^2^⟩ = 𝑁𝑏^2^, which is supported by our earlier findings that all sequences in the dense phase follow ideal chain statistics (**Figure 2 and Supplementary Figure S2**). The Rouse model predicts translational diffusivity 𝐷 as:

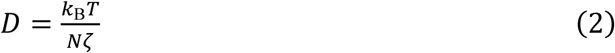

The Rouse model also yields the bulk viscosity 𝜂 of an unentangled polymer solution as:

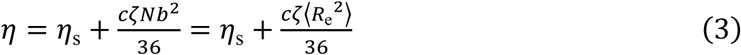

where 𝜂_s_ is the solvent viscosity, and 𝑐 is the concentration of chain segments (monomers per unit volume) in the dense phase.

We first asked whether the Rouse model could predict chain reconfiguration dynamics in our condensates using Equation (1). To test this, we computed end-to-end vector relaxation times 𝜏_Re_ in E-K condensates from the autocorrelation function 𝑐_e_ of the end-to-end vector 𝑹_e_, using the same procedure as for obtaining 𝜏_𝑝_ from the normal modes. The autocorrelation 𝑐_e_ decayed to zero for all sequences (**Supplementary Figure S10**), highlighting complete chain relaxation. The sequence-dependent changes in 𝜏_Re_ (**Figure 5b**) mirrored the dynamic slowdown observed in viscosity and diffusivity. At fixed *N*, increasing nSCD slowed chain reconfiguration consistent with enhanced electrostatic interactions. Relaxation times 𝜏_Re_ scaled with 𝑁 according to a power law, with exponents ranging from 2.44 to 2.77 across different nSCD values. As with macroscopic dynamics, these exponents fall within the crossover regime between Rouse and reptation-like dynamics^28,29,47^. We next tested Equation (1) using simulation-derived quantities to predict 𝜏_Re_: the measured ⟨𝑅_e_^2^⟩ and the Langevin friction coefficient 𝜁 applied in the simulations. The Rouse model systematically underestimated 𝜏_Re_, with errors increasing at higher nSCD values (**Figure 5c**). Predictions for diffusivity and viscosity using Equations (2) and (3) showed similarly poor agreement with simulation data (**Figure 5d and Supplementary Figure S11**). These discrepancies suggest that the effective friction experienced by monomers in the dense phase exceeds the nominal Langevin friction 𝜁 specified in the simulations.

### Chain Relaxation Times Enable Quantitative Prediction of Condensate Material Properties through Rouse Theory

Although the Rouse model does not quantitatively capture sequence-specific effects on chain relaxation times, viscosity, or diffusivity, we asked whether it could still serve as a useful framework by connecting chain-level relaxation to macroscopic properties. A recent study demonstrated that this is possible by using measured relaxation times in place of model parameters such as the monomeric friction coefficient 𝜁^21^. In this approach, the diffusivity *D* can be expressed directly in terms of the chain relaxation time 𝜏_R_:

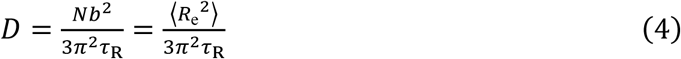

Similarly, viscosity 𝜂 can be written as a function of either 𝐷 or 𝜏_R_ as:

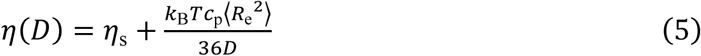

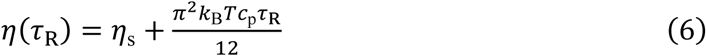

where 𝑐_p_ = 𝑐/𝑁 is the protein concentration in the dense phase.

We applied these equations using simulation-derived values of ⟨𝑅_e_^2^⟩ and 𝜏_Re_ (used in place of 𝜏_R_) and found that the Rouse model accurately predicted both 𝜂 and 𝐷 across all sequences (**Supplementary Figure S12**). Notably, this relationship holds even when solvent effects are explicitly considered using the Martini model (**Supplementary Figure S12**). In contrast, incorporating entanglement corrections into the Rouse model did not improve agreement with our simulation data (**Supplementary Figure S13**; see Methods). These results indicate that the Rouse model, even without entanglement corrections, provides a quantitative link between chain relaxation and condensate material properties in the crossover regime. This finding leads to a new question: what governs the internal chain relaxation times, such as 𝜏_Re_, that underlie macroscopic properties like diffusivity and viscosity?

As a preliminary step, we tested whether replacing 𝜁 in Equation (1) with monomer relaxation times 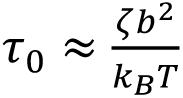 measured directly from normal model analysis of the dense phase could improve the Rouse model’s predictions. Specifically, we utilized the following relation:

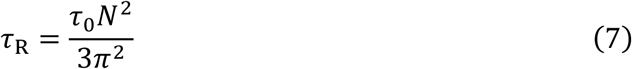

Using 𝜏_0_led to slightly better agreement between Rouse predictions and simulation results (**Supplementary Figure S14**), compared to using 𝜁. This improvement likely stems from the fact that 𝜏_0_implicitly accounts for crowding effects present in the dense phase. However, deviations persisted across different nSCD values, as 𝜏_0_ does not capture sequence-specific effects (**Figure 4c and Supplementary Figure S9**). We also tested whether incorporating entanglement corrections into the Rouse model (see Methods) could improve the prediction of 𝜏_Re_, using both 𝜁 and 𝜏_0_as inputs. In both cases, the Rouse model with entanglement failed to reproduce the observed 𝜏_Re_ values across different sequences (**Supplementary Figure S15**).

Together, these results demonstrate that Rouse theory can quantitatively predict condensate material properties when the chain relaxation times are known. However, the Rouse framework does not allow a direct quantitative connection from sequence-level information to macroscopic material properties. To address this gap, we next investigated whether an alternative theoretical framework could explain the sequence-dependent dynamic slowdown observed in our simulated condensates across multiple length scales.

### Sticky Rouse Theory Captures Sequence-Dependent Dynamics in IDP Condensates

In charge-rich IDP condensates, oppositely charged residues form transient interchain contacts, which can slow dynamics, especially in sequences with greater charge blockiness. The sticky Rouse model^48–53^ extends the classical Rouse framework by incorporating these intermittent binding and unbinding events (**Figure 6a**). In this model, the chain relaxation time is given by

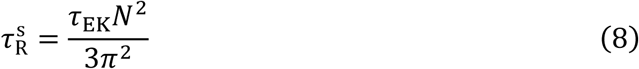

**Figure 6:**
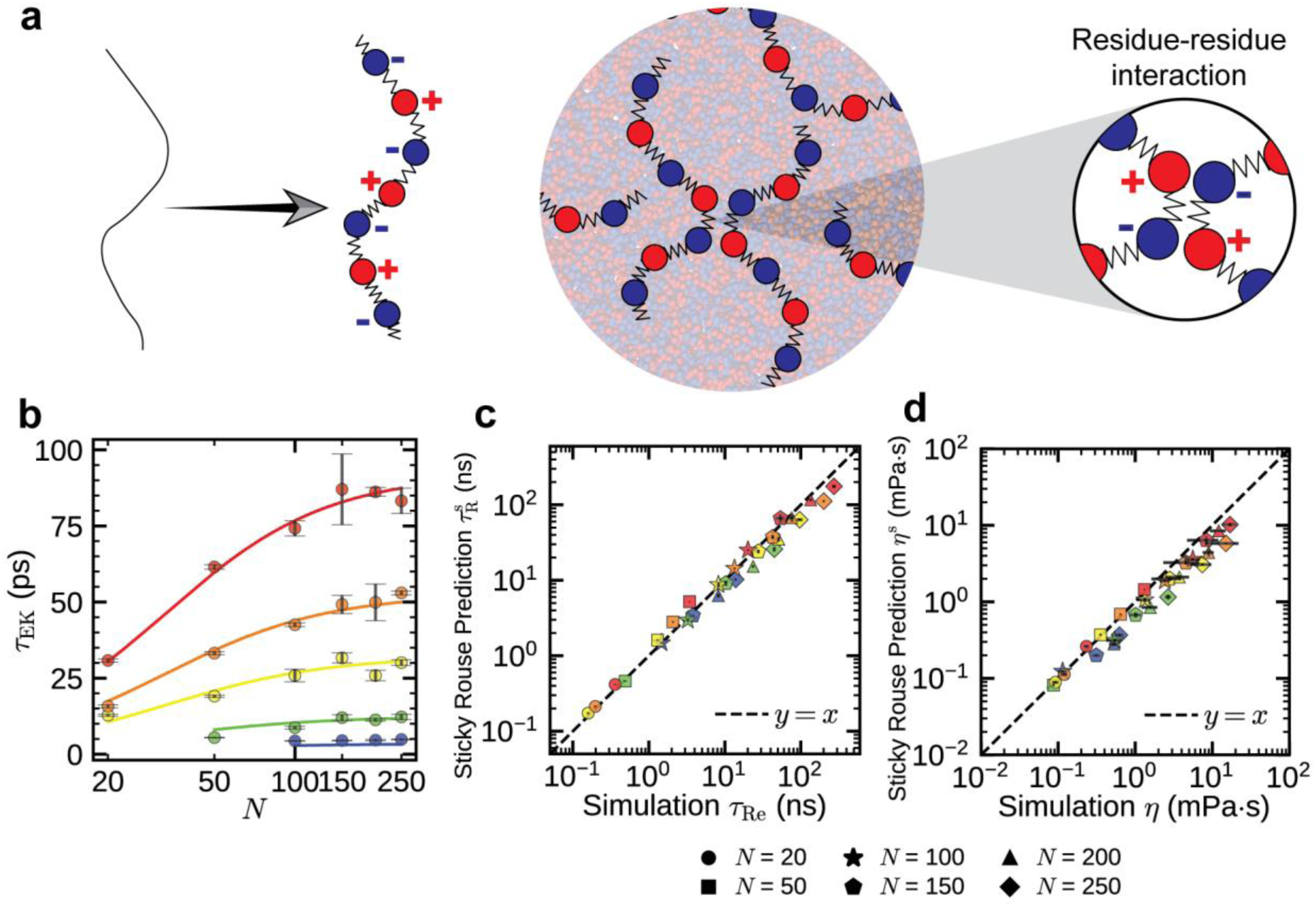
Sticky Rouse theory helps in quantitative bridging of monomer-monomer interactions with the chain relaxations as well as dense phase dynamics and rheology of IDP condensates: (a) Schematic representation of sticky Rouse model where a polymer is modeled as beads connected by springs, exhibiting random motion and slowed by sticky interactions. (b) Interchain contact lifetime 𝜏_EK_ of oppositely charged residues as a function of chain length 𝑁 and normalized sequence charge decoration nSCD within the dense phase of E-K sequences. (c) Correlation plot comparing sticky Rouse theory predictions and simulated end-to-end vector relaxation times where predictions were made using Equation (8) (d) Correlation plot comparing sticky Rouse theory predictions and simulated dense phase viscosities where predictions were made by substituting predicted chain reconfiguration times 𝜏_R_^s^ in Equation (6). The solid lines in (b) represent the phenomenological model fits, while dashed line in (c) and (d) corresponds to the 𝑦 = 𝑥 line. Symbol colors, ranging from blue to red, indicate increasing nSCD values.

where 𝜏_EK_ is the average contact lifetime between oppositely charged residues, replacing 𝜏_0_ in Equation (7) as the relevant monomer relaxation timescale.

To characterize contact dynamics in our simulations, we quantified 𝜏_EK_ for each sequence, which reflects how long electrostatic contacts persist, including intermittent breaking and reformation^22,25,26^ (see Methods). We found that 𝜏_EK_ increased with chain length 𝑁 and eventually plateaued (**Figure 6b**). For sequences with nSCD ≤ 0.20, the plateau occurred beyond 𝑁 = 100, while for nSCD > 0.20, it occurred beyond 𝑁 = 150, even though macroscopic properties continued to show dynamic slowdown. At fixed 𝑁, increasing nSCD led to longer-lived contacts, consistent with stronger electrostatic interactions between charged residues. This contrasts with the intrachain monomer relaxation time 𝜏_0_, which did not capture this sequence dependence, highlighting 𝜏_EK_ as a key sequence-sensitive timescale.

To better understand how sequence features influence 𝜏_EK_, we developed a phenomenological model based on the sticky Rouse framework (see Methods). The model incorporates the effects of charge patterning and viscosity, and it allows 𝜏_EK_ to be estimated directly from sequence features such as *N* and nSCD. The resulting expression accurately captures the simulation trends across all sequences (solid lines in **Figure 6b**) and provides a direct link between sequence-level features and local interaction timescales.

We then asked whether τ_EK_, when used as input in the sticky Rouse model, could quantitatively predict the chain relaxation times 𝜏_Re_ observed in simulations. Indeed, the predicted 𝜏^s^ values matched simulation-derived 𝜏_Re_ across all sequences, with the data collapsing onto the identity line (**Figure 6c**). Because earlier analysis established that 𝜏_Re_ is directly coupled to macroscopic observables, this accurate prediction of 𝜏_Re_ implies that diffusivity and viscosity can also be recovered via τ_EK_. Indeed, using 𝜏_R_^s^ in the corresponding Rouse expressions, we found that 𝐷 and 𝜂 can be quantitatively predicted across the sequence sets investigated (**Figure 6d and Supplementary Figure S16**). To test whether entanglement effects might further refine these predictions, we also applied a sticky Rouse model with reptation-like corrections. While the extension preserved the nSCD-dependent trend, seen in the collapse of data across sequences, it consistently overestimated 𝜏_Re_ (**Supplementary Figure S17**; see Methods). This suggests that entanglement dynamics are not the dominant contributor in these condensates. Together, these results demonstrate that the sticky Rouse framework, when paired with a sequence-informed model of contact lifetimes, provides a robust and quantitative connection from monomer-level interactions to chain relaxation and ultimately to the macroscopic material properties of IDP condensates.

## DISCUSSION

Our findings offer a generalizable framework for predicting how sequence-level features of intrinsically disordered proteins (IDPs) govern the material properties of biomolecular condensates. By anchoring the analysis in chain relaxation times, we establish a mechanistic link between residue-level interactions and emergent material properties. Through systematic variation of chain length *N* and charge patterning nSCD, we demonstrate that longer and more charge-segregated sequences give rise to slower dynamics in the dense phase. This slowdown is better captured by chain-level relaxation times than by monomer-level friction, highlighting the importance of cooperative effects in IDP-rich condensates. The observed scaling of relaxation times and material properties with *N* places our systems in a crossover regime between Rouse and reptation dynamics, suggesting that entanglements are emerging but not yet dominant. Notably, the chain lengths examined here are comparable to those found in biologically relevant IDPs^30^, implying that the crossover behavior we observe may be broadly applicable to natural systems.

While the classical Rouse theory models a polymer as a series of beads connected by springs with no specific interactions^21,28,29^, this simplification limits its predictive power for complex biological systems where residue-level interactions, particularly electrostatics^54,55^, are key. In our simulations, although the chains exhibit ideal conformational statistics, the dynamic behavior is dominated by sequence-specific electrostatic interactions. These interactions lead to a systematic breakdown of the Rouse predictions when material properties are computed using the nominal monomeric friction. By contrast, the sticky Rouse model incorporates the effect of transient inter-residue interactions and allows us to replace monomer relaxation times with contact lifetimes 𝜏_EK_, enabling accurate prediction of both chain and condensate-level dynamics. Importantly, our phenomenological model relates 𝜏_EK_ directly to sequence features such as nSCD, allowing the full sequence-dynamics-material property link to be quantitatively established.

Our study focuses on charge-neutral E–K sequences, which allow controlled tuning of charge patterning while avoiding confounding effects from net charge or hydrophobicity. These findings are directly relevant to many natural IDPs that exhibit varied charge distributions^56,57^, and they provide a baseline for future studies incorporating additional sequence complexity. For example, the influence of hydrophobic or aromatic residues, which can contribute^58–61^ to short-range and directional interactions, can be explored in future extensions of the sticky Rouse framework. Overall, this work lays the foundation for predictive sequence-to-property design principles for biomolecular condensates with tailored material characteristics.

## METHODS

### Sequence Generation

To investigate the effects of chain length 𝑁 and charge patterning on the material properties of dense phases, we utilized charge-neutral polyampholytes composed of equal numbers of glutamic acid (denoted as ‘E’, net charge 𝑞_E_ =-e, diameter 𝜎_E_ = 5.92 Å, molar mass 𝑚_E_ = 129.1 g/mol) and lysine (denoted as ‘K’, net charge 𝑞_K_ = +e, diameter 𝜎_K_ = 6.36 Å, molar mass 𝑚_K_ = 128.2 g/mol). These sequences served as model intrinsically disordered proteins (IDPs) with 𝑁 values ranging from 20 to 250. For each 𝑁, we systematically varied the charge patterning, creating sequences that ranged from perfectly alternating arrangements of E and K to fully segregated blocks of E and K. To quantify the distribution of charges along the chain, we used the sequence charge decoration (SCD) parameter^31,32^:

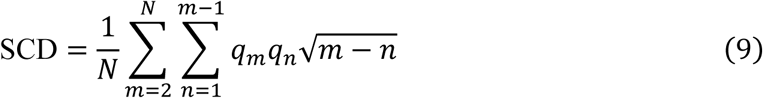

In this representation, sequences with alternating E and K residues exhibit the maximum SCD value (SCD_max_), while sequences with a diblock distribution have the minimum SCD value (SCD_min_). To compare sequences of different charge patterning and lengths, we normalized the SCD values using the minimum and maximum SCD limits for each 𝑁, resulting in the normalized SCD (nSCD) defined as:

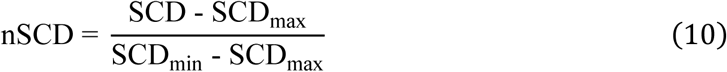

With this normalization, an nSCD of 0 corresponds to a perfectly alternating sequence, while an nSCD of 1 corresponds to a diblock sequence, regardless of the length of the sequence^36,62^. We selected sequences with nSCD values of 0, 0.02, 0.20, 0.46, and 1 for each 𝑁, allowing a systematic investigation of the impact of charge patterning on the dense phase material properties. E-K sequences generated are shown in **Supplementary Table S1**.

### Hydropathy Scale (HPS) Model for IDPs

To model the IDP sequences, we used a recently developed coarse-grained approach where each residue is represented by a single bead^63,64^. Bonded interactions between residues were modeled with a harmonic potential:

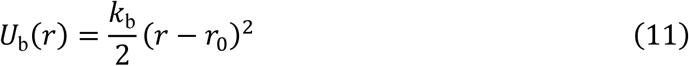

In this equation, 𝑟 is the distance between bonded residues with 𝑘_b_ = 20 kcal/(mol Å^2^) as the bond spring constant and 𝑟_0_= 3.8 Å as the equilibrium bond length. For non-bonded residues, we applied a modified Lennard-Jones potential 𝑈_vdW_(𝑟), which allows the attractive interactions to be scaled based on the average hydropathy 𝜆 = ^𝜆𝑖^<u>^+^</u>^𝜆^^𝑗^ of residues 𝑖 and 𝑗^65,66^:

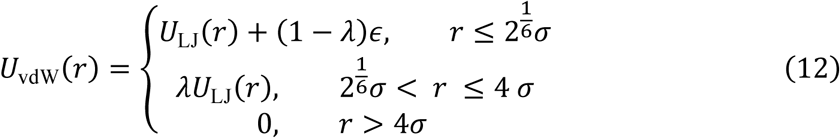

The standard Lennard-Jones potential 𝑈_LJ_(𝑟) is expressed as:

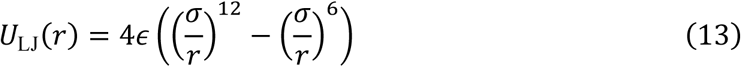

Here, 𝜎 corresponds to the average diameter of the interacting residues and 𝜖 = 0.2 kcal/mol reflects the interaction strength. For E-K sequences, the 𝜆 values were determined using the Kapcha-Rossky scale^64,67^. Charged residues interact via a Coulomb potential 𝑈_𝑒_(𝑟) with Debye-Hückel electrostatic screening^68^:

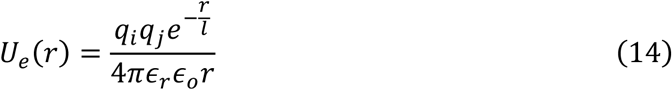

Here, 𝜖_𝑜_ represents the permittivity of free space, 𝜖_𝑟_ = 80 denotes the relative permittivity, and 𝑙 = 10 Å is the Debye screening length, representative of an aqueous environment typical of physiological conditions, with an electrostatic cut-off of 35 Å.

### Simulation Details

To study the dense phase of E-K sequences with varying 𝑁 and nSCD, we conducted NPT simulations of the sequences in a cubic simulation box at a constant pressure of 0 atm for a total duration of 0.5 𝜇s. During this time, the sequences reached their preferred dense phase concentration 𝜌 (**Supplementary Figure S18**). Following this, we transitioned to Langevin dynamics (LD) simulations in the NVT ensemble, adjusting the simulation duration based on 𝑁 and nSCD. Sequences with longer chain lengths and higher charge segregation, particularly those with 𝑁 = 250 and nSCD = 0.02, 0.20, 0.46, and 1, as well as 𝑁 = 200 with nSCD = 0.20, 0.46, and 1, required longer simulation runs of 2.5𝜇s to capture the diffusive motion of chains. For the remaining E-K sequences, NVT simulations were performed for 1𝜇s. All dense-phase NVT simulations included a 40-ns equilibration period, and only the data after this equilibration period were used for analysis. Notably, E-K sequences of 𝑁 = 20 and 50 with nSCD = 0, as well as 𝑁 = 20 with nSCD = 0.02, did not form a stable dense phase during the P = 0 atm simulation. This suggests that their perfectly alternating charge arrangement led to weaker intermolecular attractions^69^. The 𝑁 = 50 trajectories from our recently published work^22^ were used for all the analyses involving 𝑁 = 50 in this work.

Single-chain simulations of E-K sequences of varying 𝑁 were conducted in a cubic box with an edge length of 400 Å, ensuring that the box size was large enough to avoid artificial self-interactions between the chains and their periodic images. Each simulation was run for a total duration of 1𝜇s.

All simulations were performed using HOOMD-blue (version 2.9.3)^70^ with features extended using azplugins (version 0.10.1)^71^. Dense phase simulations were conducted for 1500, 600, 300, 200, 150, and 120 chains of E-K sequences for 𝑁 = 20, 50, 100, 150, 200, and 250, respectively. Simulations were conducted at 300 K with periodic boundary conditions applied in all three Cartesian directions with a timestep of 10 fs and a Langevin damping factor of 𝑡_damp_ = 1000 ps was applied to set the friction coefficient 𝜁_𝑖_ of a residue of mass 𝑚_𝑖_ in the chain to 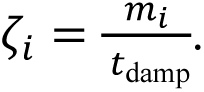 Trajectories were saved at intervals ranging from 0.1 ps to 500 ps to compute segmental dynamics, contact dynamics, and the condensate material properties.

### Calculation of Dense Phase Viscosity and Solvent Viscosity

To determine the zero-shear bulk viscosity of the dense phase, we used the Green-Kubo relation^39–41,72–77^. First, we used the autocorrelation of the off-diagonal components of the stress tensor (computed on-the-fly using HOOMD’s autocorrelator tool^78,79^) to compute shear stress relaxation modulus 𝐺(𝑡) in the limit of zero deformation as

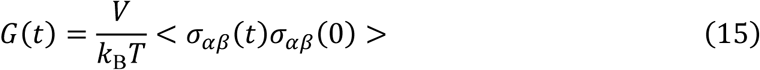

where 𝜎_𝛼𝛽_ is an off-diagonal component (𝛼𝛽) of the stress tensor, 𝑉 is the volume, 𝑘_B_ is the Boltzmann constant, and 𝑇 is the temperature. At equilibrium, 𝐺(𝑡) is then obtained using three independent off-diagonal components of the stress tensor (**Supplementary Figure S19**) as

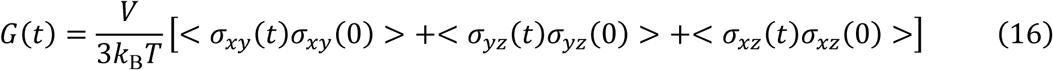

To mitigate the noise in the shear stress relaxation modulus 𝐺(𝑡) at longer times, we averaged 𝐺(𝑡) over several independent NVT simulations, following the methodology discussed by Zhang et al^75^. Finally, we estimated the zero-shear viscosity 𝜂(𝑡) by directly integrating the 𝐺(𝑡) using the Green–Kubo relation

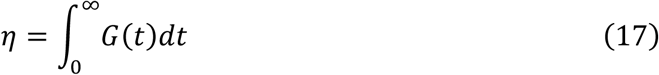

and computed average viscosities from the independent simulations conducted for different *N* and nSCD values (**Figure 3b and Supplementary Figure S20**). The duration of the NVT simulations was chosen based on the chain length and charge patterning of the sequences. Longer and more charge-segregated chains required extended simulation times to achieve convergence in 𝜂 values. Detailed run durations and number of replicas for each sequence are provided below.

To calculate the solvent viscosity 𝜂_s_ (**Supplementary Figure S21c**), we utilized the Stokes-Einstein equation:

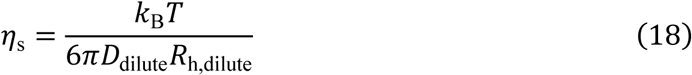

where 𝐷_dilute_ is the diffusion coefficient of the chain in the dilute phase (**Supplementary Figure S21b**), and 𝑅_h,dilute_ is the hydrodynamic radius of the chain in the dilute phase (**Supplementary Figure S21a**). The hydrodynamic radius was calculated using the Kirkwood approximation:

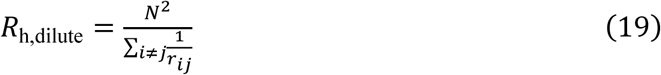

where 𝑟_𝑖𝑗_ is the distance between atoms 𝑖 and 𝑗.

### Normal Mode Analysis

We computed the normal modes 𝑿_𝑝_ of E-K sequences inside the dense phase as^36,42–45^

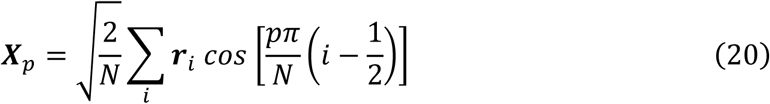

Here, 𝒓_𝑖_ is the position vector of the 𝑖-th monomer and mode number 𝑝 corresponds to a subchain of 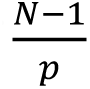 segments for a chain of length 𝑁. Therefore, modes 1 ≤ 𝑝 ≤ 𝑁 − 1 describe the internal relaxation of a chain. Specifically, 𝑝 = 1 corresponds to the slowest chain-level dynamics. The autocorrelation function 𝑐_𝑝_ of 𝑿_𝑝_ (**Supplementary Figure S7**), which is well described by a stretched exponential function 𝑐_𝑝_(𝑡) = 𝑒^−(𝑡/𝜏)𝛽^, is given by

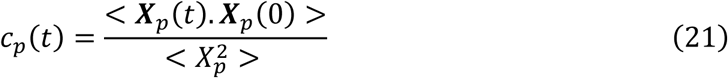

where 𝜏 is the characteristic time scale, and 𝛽 is the stretching exponent.

The characteristic relaxation time 𝜏_𝑝_ of mode 𝑝 (**Supplementary Figure S8**) is calculated as follows:

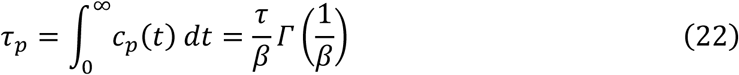

Note that 𝛤 is the gamma function.

### Calculation of Monomer-Monomer Lifetimes

To estimate contact lifetimes of oppositely charged residues in the dense phase, we computed the intermittent contact time autocorrelation function^22,25,26^ 𝑐(𝑡) as (**Supplementary Figure S22**):

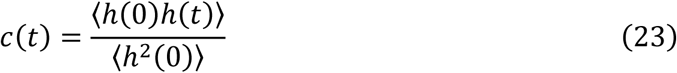

where ℎ(𝑡) is a step function that equals 1 if a pair of oppositely charged residues 𝑖 and 𝑗 are in contact at time 𝑡, and 0 otherwise. A contact was defined as occurring when the distance between residues was below a cut-off of 1.5𝜎, where 𝜎 is the average diameter of E and K residues. The autocorrelation function captures how long contacts persist, including repeated breaking and reforming of interactions. The characteristic contact lifetime 𝜏_EK_ was determined by integrating 𝑐(𝑡) up to the time at which it decayed to 0.05^22,24^.

### Phenomenological model for sequence-dependent contact lifetimes

To quantitatively model the effect of sequence on the contact lifetime τ_EK_, we built on the sticky Rouse framework^49,53^, where the sticker lifetime is given by:

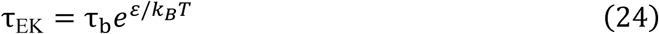

Here τ_b_ is the baseline contact time in the absence of specific bonding interactions (primarily due to random collisions), and 𝜀 is an effective interaction energy between oppositely charged residues. To account for sequence effects, we decomposed 𝜀 into:

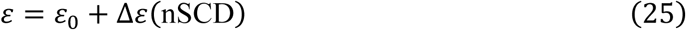

where 𝜀_0_is contact energy for a perfectly alternating sequence, and Δε captures the influence of charge clustering. We estimated Δε ≈ α𝑛_𝑐_, where 𝑛_𝑐_ is the average number of like-charge nearest neighbors, and found empirically that 𝑛_𝑐_ = 1.76 nSCD^0.24^ (**Supplementary Figure S23**).

We also accounted for viscosity effects on τ_b_, using:

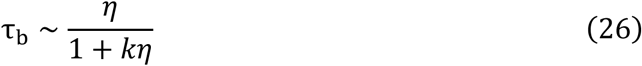

where 𝜂 = 𝑎𝑁^𝛾^ and 𝑘 is a constant. Combining these expressions, we obtained the following empirical equation:

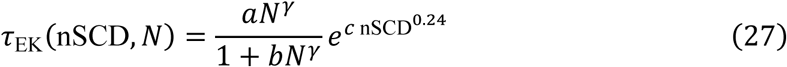

Where the fitted parameters were γ = 1.44, 𝑎 = 0.022, 𝑏 = 0.0066, and 𝑐 = 3.2. These values were selected to best fit the trends in simulation data across all sequence variants. The scaling exponent γ closely matched the chain length dependence of viscosity, supporting the idea that longer chains hinder contact dynamics.

### Rouse Theory with Entanglement Corrections

To evaluate whether chain entanglements contribute to the dynamics of IDPs in our simulated dense phases, we tested an extended version of the Rouse model that includes entanglement corrections. In this framework^21,29^, Rouse chain motion is constrained within an effective tube of diameter 𝑎 (also known as entanglement spacing), representing the spatial confinement due to surrounding chains. The chain diffusion coefficient is given by

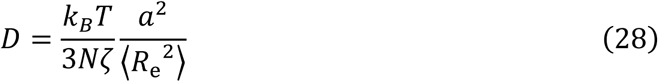

On the other hand, the dense phase viscosity of the polymer solution is given by

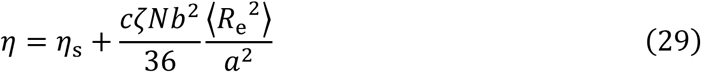

The chain relaxation time that governs the reptation dynamics is the disentanglement time τ_d_ (also known as reptation time)^21,80^, which is the time it takes for a chain to disentangle from a tube within which it was confined and is given by

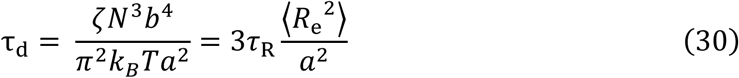

where 𝜏_R_ is the Rouse time and 𝑏 is computed using the mean squared end-to-end distance ⟨𝑅_e_^2^⟩ of chains within the dense phase using the relation ⟨𝑅_e_^2^⟩ = 𝑁𝑏^2^. τ_d_ can also be written in terms of the monomer relaxation times 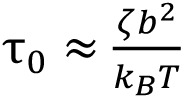 as

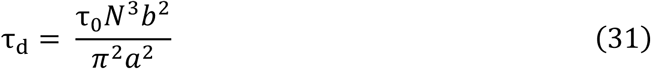

On the other hand, we can also express the dense phase properties in terms of the chain disentanglement time. By combining Equations (28) and (30), 𝐷 can be written in terms of 𝜏_d_ as:

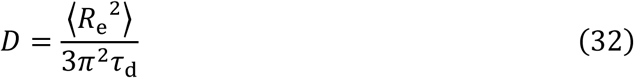

This expression matches the form seen in the original Rouse model (Equation (4)), indicating that reptation theory does not introduce additional correction terms for predicting the diffusion coefficient from chain relaxation dynamics. Therefore, we did not separately compute 𝐷 using the simulation-derived 𝜏_Re_ in the entanglement-corrected Rouse equation, as it would yield the same result as the standard Rouse expression 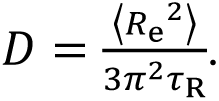.

Similarly, combining Equations (29) and (30), 𝜂 can be expressed in terms of 𝜏_d_ as:

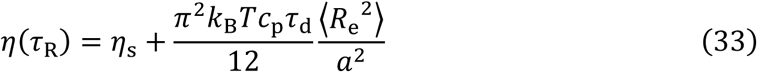

where 𝑐_p_ (= 𝑐/𝑁) is the protein concentration in the dense phase. First, we estimated the entanglement spacing 𝑎 for E-K sequences in our simulated condensates through topological analysis using the 𝑍1+ algorithm developed by Kröger’s group^81,82^. This method constructs primitive paths by minimizing the chain’s contour lengths under the constraint of fixed ends. From the resulting primitive path configuration, the mean number of entanglements (also referred to as kinks) per chain ⟨𝑍⟩ was estimated (**Supplementary Figure S24a**). The entanglement length 𝑁_e_^M−kink^, defined as the number of monomers between successive kinks, was then estimated (**Supplementary Figure S24b**) as 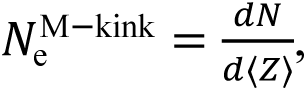 where 𝑁 is the chain length. The corresponding entanglement spacing 𝑎 (**Supplementary Figure S24c**) was computed using the relation^80^:

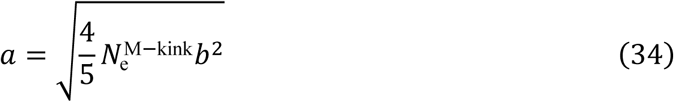

We used simulation-derived quantities ⟨𝑅_e_^2^⟩, 𝜏_0_, and 𝜏_Re_ (as 𝜏_d_), along with 𝑎 from topology analysis and the mean monomer friction (𝜁), in Equations (30,31,33) to predict 𝜂 and 𝜏_Re_

### (Supplementary Figures S13 and S15).

We also evaluated an extended framework by incorporating sticky interactions into the entanglement-corrected Rouse theory. Here we substituted 𝜏_0_in Equation (31) with 𝜏_EK_. In this case, the chain relaxation time becomes:

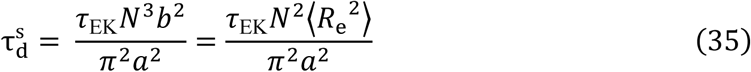

We used simulation-derived 𝜏_EK_ along with 𝑎 from topology analysis in the above equation to predict 𝜏_Re_ (**Supplementary Figure S17**).

While theoretical predictions rely on end-to-end vector relaxation times, we also computed end-to-end distance relaxation times and found them to be about three times shorter than vector relaxation times (**Supplementary Figure S25**). This provides a quantitative reference for interpreting experimental measurements, which typically has access to only distance-based relaxation in Förster resonance energy transfer (FRET) experiments. The shorter distance relaxation times observed here could contribute to the apparent necessity of including entanglement corrections in experimental analyses^21^, as the measurements are based on distance relaxation rather than vector relaxation.

### Martini Simulations

The Martini data shown in **Supplementary Figure S12** are from trajectories generated in our recently published work^22^ and for the sake of completeness, we briefly summarize the simulation details. We simulated E-K sequences (𝑁 = 50) using the Martini 3 coarse-grained force field with explicit solvent and ions^83^. The protein chains were coarse-grained using martinize2 Python script, and non-bonded interactions were modeled using the Verlet cut-off scheme with a 11 Å cut-off for van der Waals and electrostatic interactions. Long-range electrostatic interactions were treated using the reaction-field method with a dielectric constant of 15^84^.

To generate the dense phase, we initially placed 80 chains in a slab geometry (100 Å × 100 Å × 740 Å) and solvated them with ∼50,000 water particles using the insane python script^85^. After energy minimization with the steepest descent algorithm, we equilibrated the system in the NPT ensemble (T = 300 K, P = 1 bar along the z-direction) for 20 ns using a 20 fs timestep. The production run was conducted for 18 μs under the same conditions, allowing the slab to reach equilibrium. The dense phase was then extracted and simulated in a cuboid box at 100 mM salt concentration to match HPS model conditions.

To compute the translational diffusivity 𝐷, we performed NVT simulations at 300 K for 10 μs and extracted 𝐷 values from the long-time regime where MSD scaled linearly with time. To compute the zero-shear viscosity 𝜂 using the Green-Kubo relation^40,72–77^, we conducted 10 replica simulations, each lasting 10 μs, and averaged 𝜂 over all replicas. All simulations were performed using GROMACS (version 2023.1)^86^ with a velocity-rescaling thermostat^87^ (time constant of 1 ps) and a Parrinello–Rahman barostat^88^ (time constant of 12 ps). Mean end-to-end distances and end-to-end vector relaxation times were calculated using the same procedure as described for the HPS model.

## ACKNOWLEDGMENTS

This article is based on the work supported by the National Institute of General Medical Sciences of the National Institutes of Health through grant R35GM153388. Y.C.K. acknowledges support from the U.S. Naval Research Laboratory via the Office of Naval Research Base Program. We also gratefully acknowledge the Texas A&M High Performance Research Computing (HPRC) facility, whose resources were instrumental in carrying out the simulations reported in this study.

## DATA AVAILABILITY

All study data are included in the manuscript and/or supporting information.

## AUTHOR CONTRIBUTIONS

K.M. and J.M. conceived and designed the research. K.M. performed the simulations and primary data generation. K.M. and J.M. designed the analyses. K.M., D.S.D., Y.C.K., and J.M. analyzed the data. K.M. wrote the original manuscript draft, and all authors contributed to the manuscript review and editing.

## Supplementary Information

**Figure S1:**
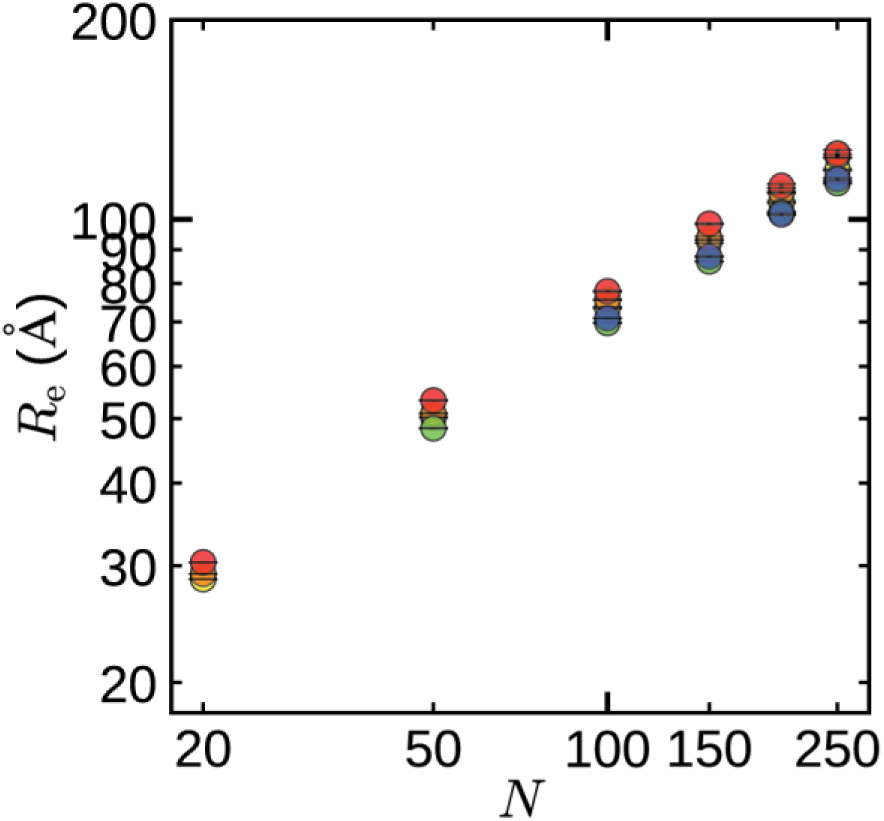
Mean End-to-end Distance Within the Dense Phase: Mean end-to-end distance 𝑅_e_ of E-K sequences as a function of chain length 𝑁 and normalized sequence charge decoration nSCD value within the dense phase. Symbol colors, ranging from blue to red, indicate increasing nSCD values.

**Figure S2:**
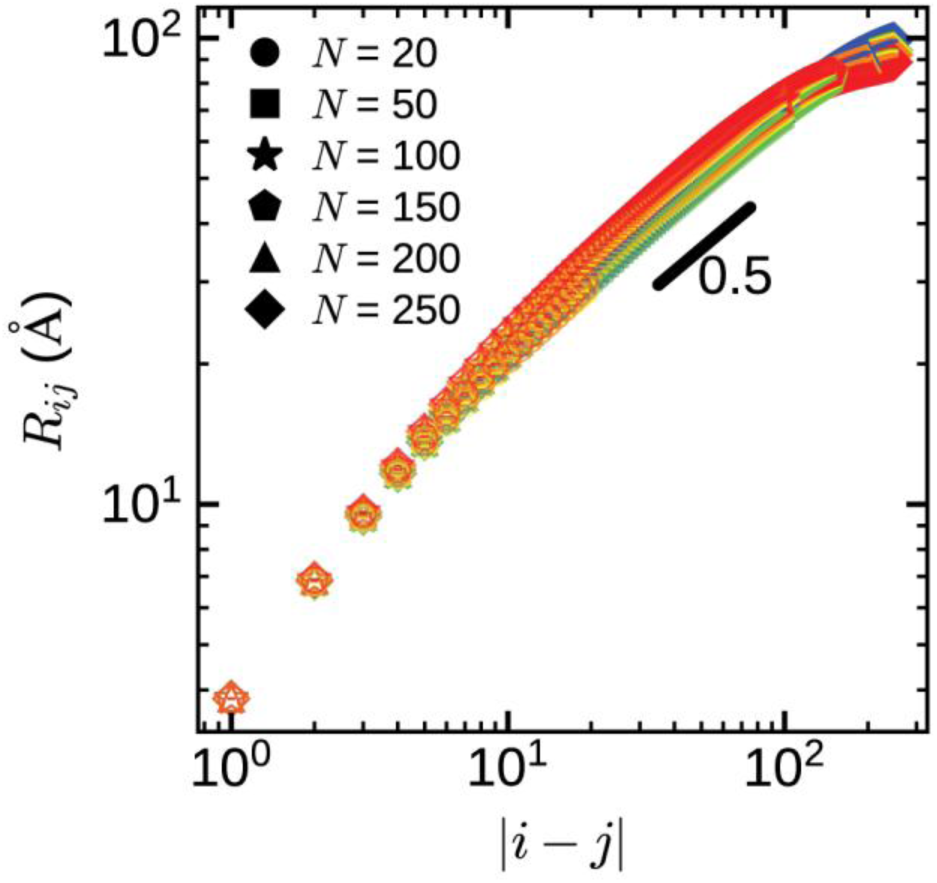
Average Interresidue Distance Within the Dense Phase: Interresidue distance 𝑅_𝑖𝑗_ of E-K sequences as a function of residue separation |𝑖 − 𝑗|, chain length 𝑁, and normalized sequence charge decoration nSCD value. Symbol colors, ranging from blue to red, indicate increasing nSCD values.

**Figure S3:**
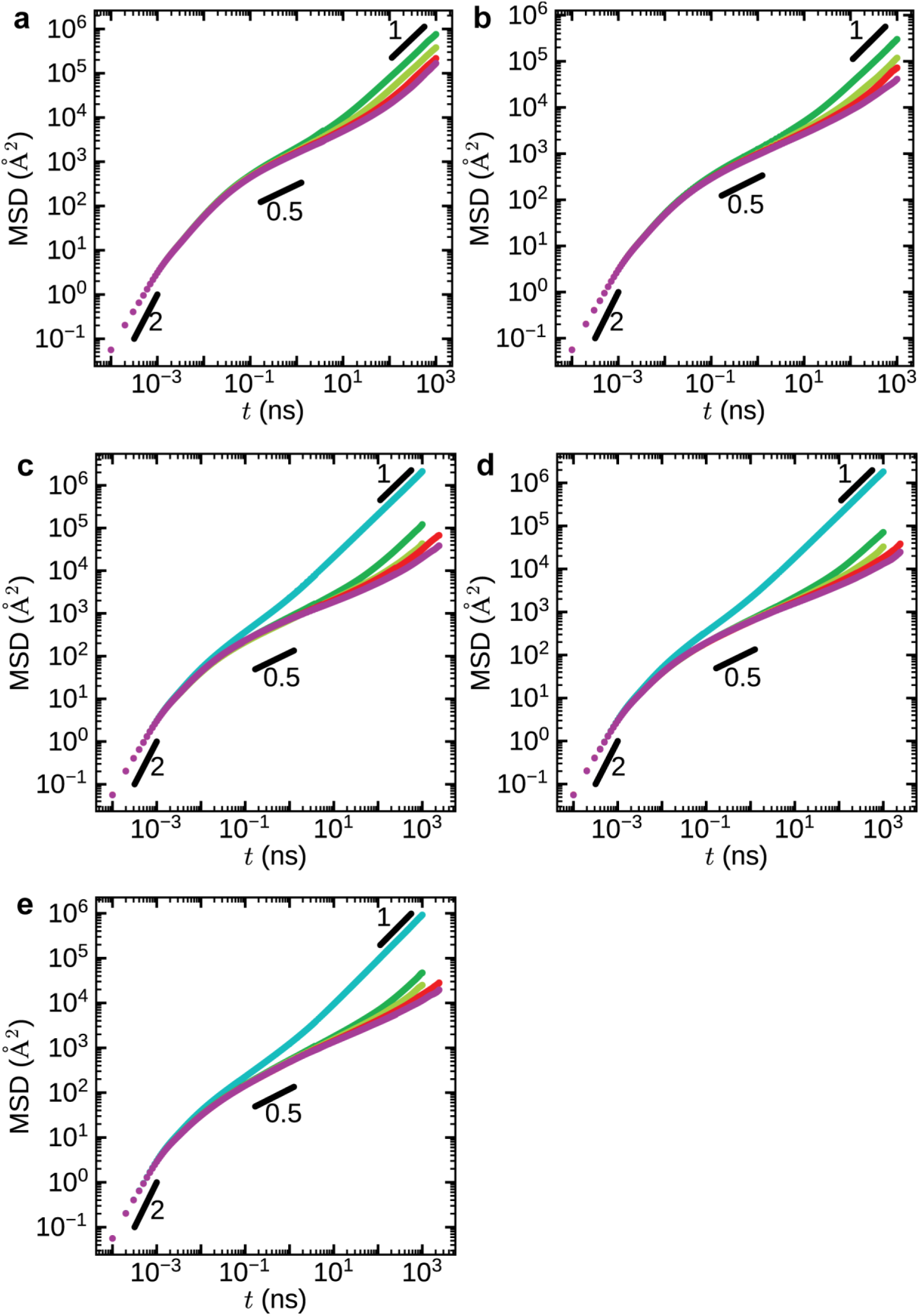
Mean Square Displacement of Residues in E-K Sequences Within the Dense Phase: Mean square displacement MSD of residues in E-K sequences as a function of time 𝑡 and chain length 𝑁 for different normalized sequence charge decoration nSCD values: (a) nSCD = 0, (b) nSCD = 0.02, (c) nSCD = 0.20, (d) nSCD = 0.46, and (e) nSCD = 1. Colors, ranging from cyan to magenta, represent increasing 𝑁.

**Figure S4:**
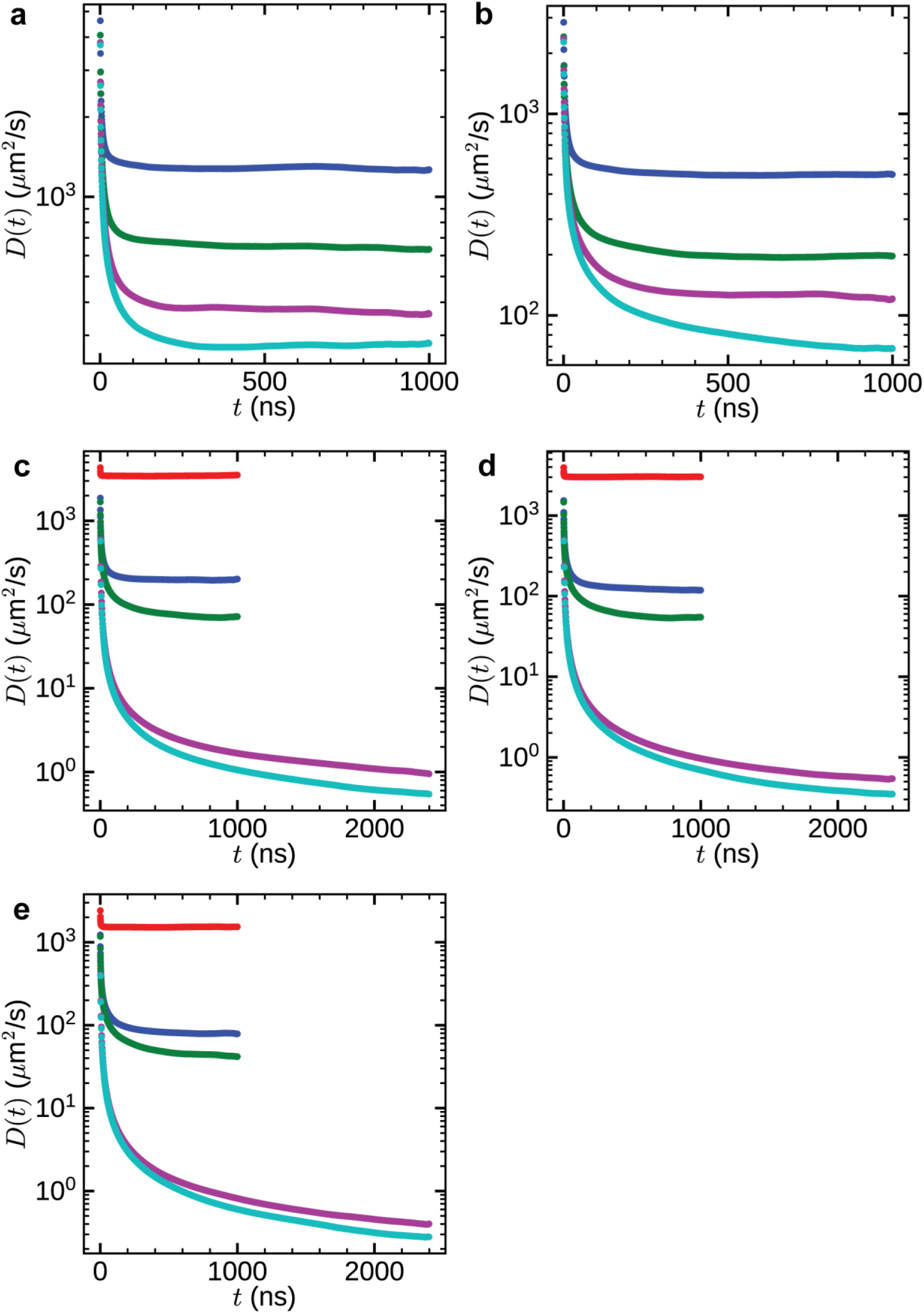
Chan Diffusion Coefficient of E-K Sequences Within the Dense Phase: Chain diffusion coefficient 𝐷 of E-K sequences as a function of time 𝑡 and chain length 𝑁 for different normalized sequence charge decoration nSCD values: (a) nSCD = 0, (b) nSCD = 0.02, (c) nSCD = 0.20, (d) nSCD = 0.46, and (e) nSCD = 1. Colors, ranging from cyan to magenta, represent increasing 𝑁.

**Figure S5:**
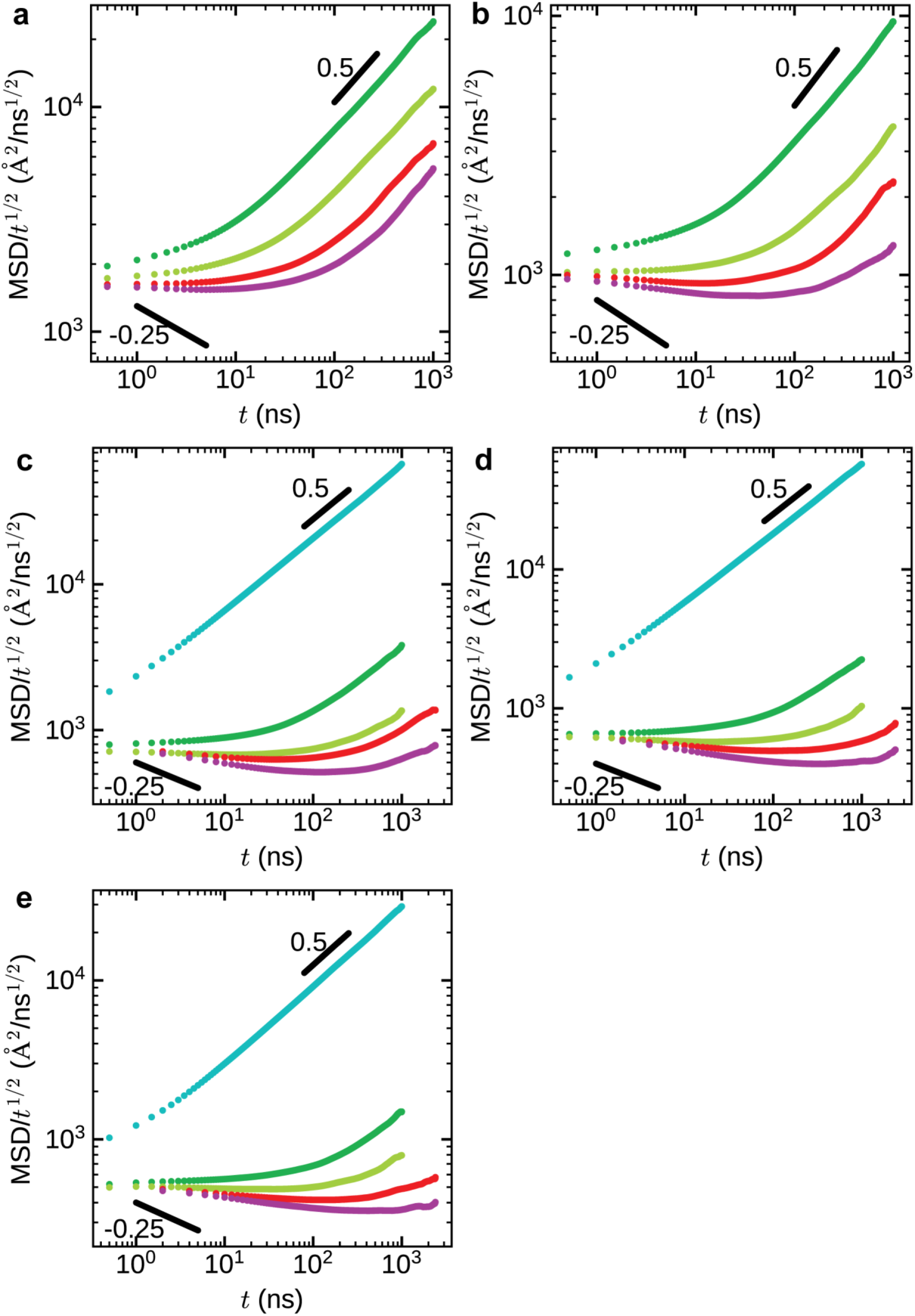
Scaled Mean Square Displacement of Residues in E-K Sequences Within the Dense Phase: Scaled mean square displacement MSD/t^1/2^ of residues in E-K sequences as a function of chain length 𝑁 for different normalized sequence charge decoration nSCD values: (a) nSCD = 0, (b) nSCD = 0.02, (c) nSCD = 0.20, (d) nSCD = 0.46, and (e) nSCD = 1. Colors, ranging from cyan to magenta, represent increasing 𝑁.

**Figure S6:**
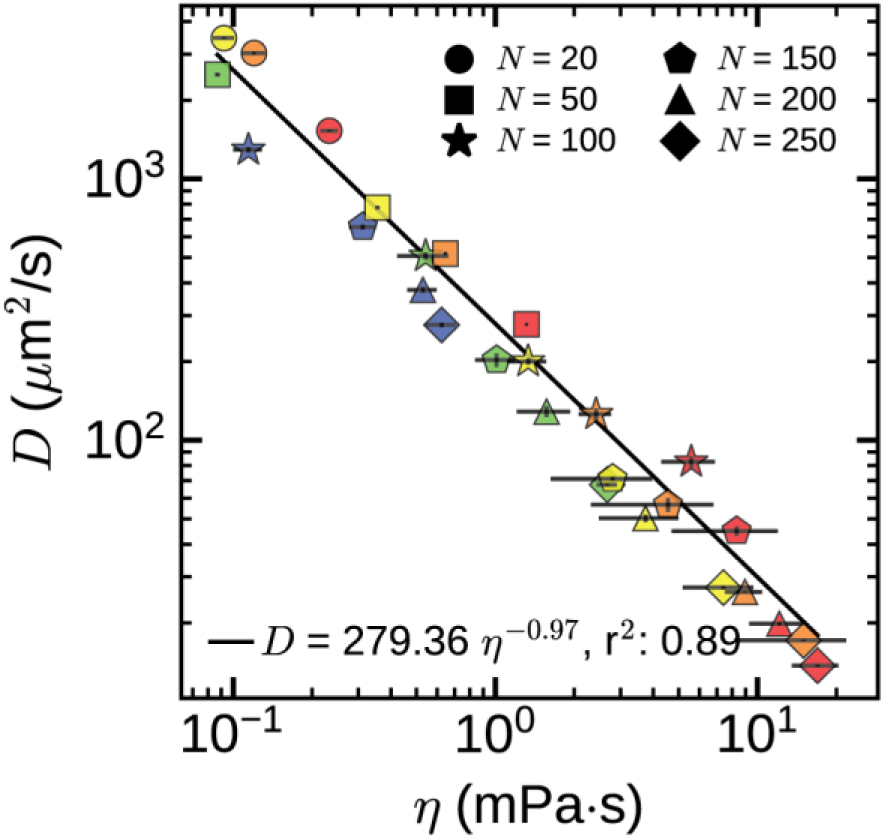
Correlation Between Diffusivity and Viscosity Within the Dense Phase: Correlation plot showing that diffusivity 𝐷 and viscosity 𝜂 obey the Stokes-Einstein equation, 𝐷 ∼ 𝜂^−1^. The solid line represents the power law fit. Symbol colors, ranging from blue to red, indicate increasing nSCD value.

**Figure S7:**
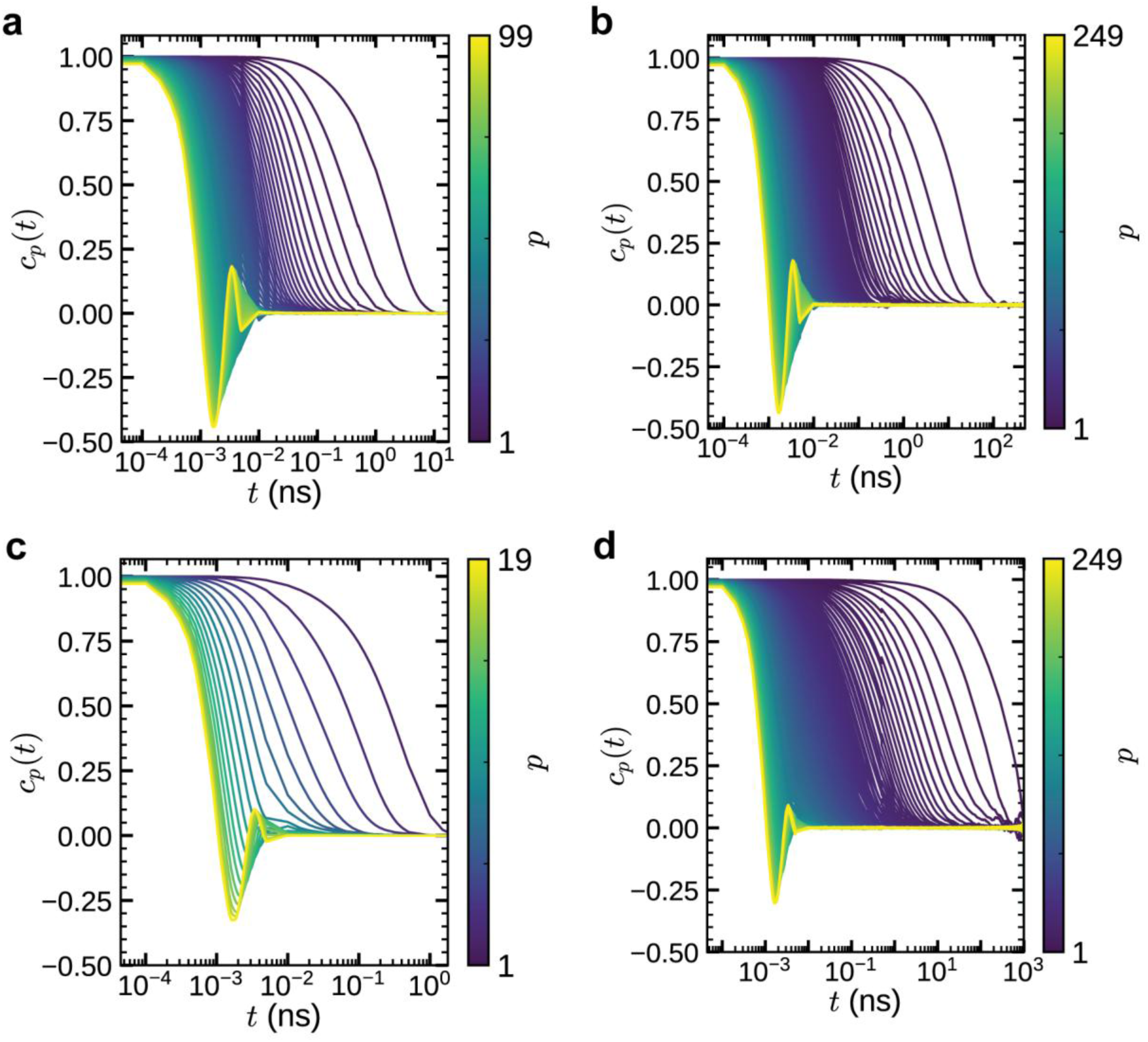
Normal Modes Autocorrelation Function for E-K Sequences Within the Dense Phase: Normal modes autocorrelation function 𝑐_𝑝_(𝑡) for modes 𝑝 = 1, 2,…, 𝑁−1 of E-K sequences for selected extreme chain lengths 𝑁 and extreme normalized sequence charge decoration nSCD values within the dense phase: (a) nSCD = 0 (𝑁 = 100), (b) nSCD = 0 (𝑁 = 250), (c) nSCD = 1 (𝑁 = 20), and (d) nSCD = 1 (𝑁 = 250).

**Figure S8:**
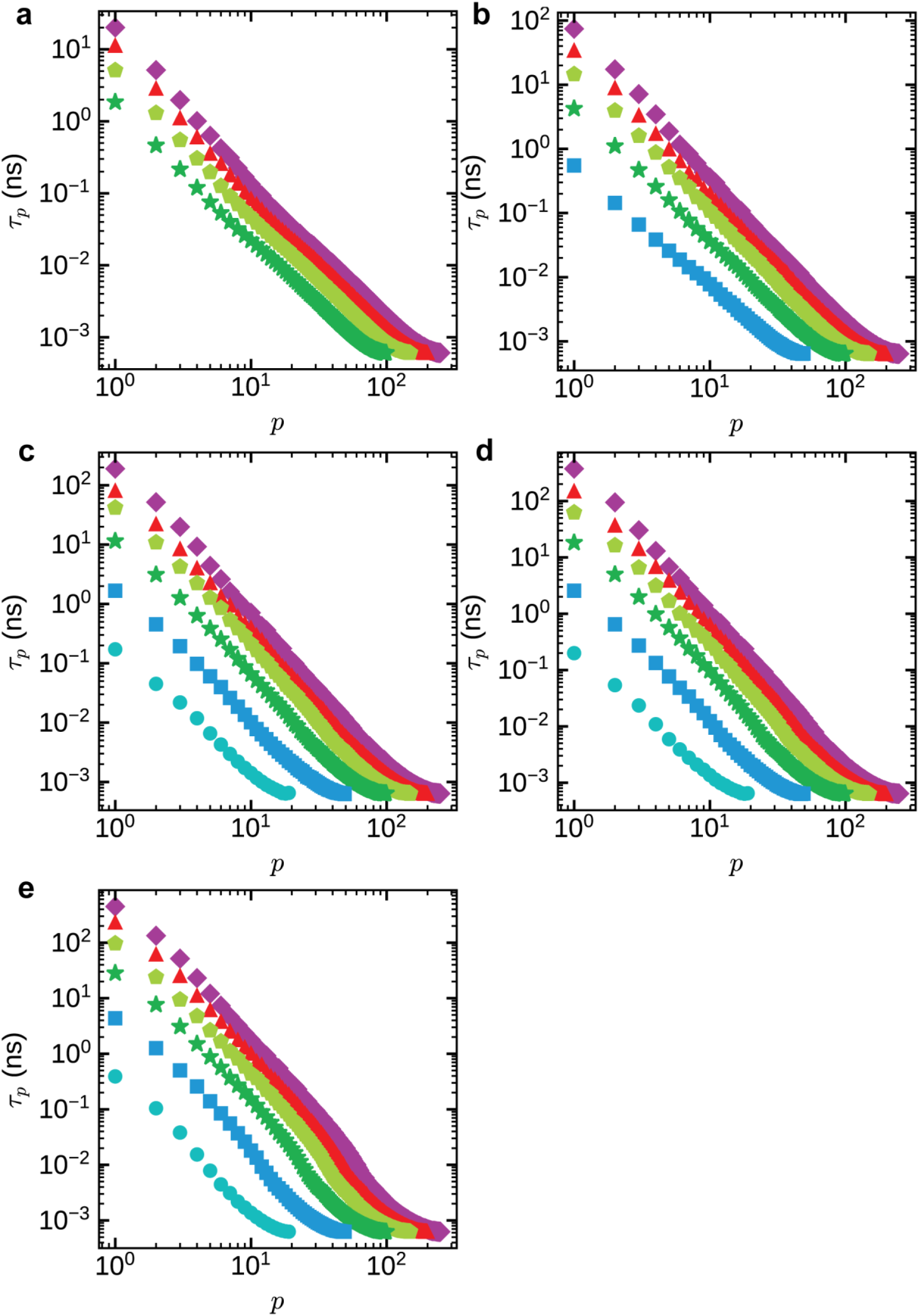
Relaxation Time of Normal Modes Autocorrelation for E-K Sequences Within the Dense Phase: Relaxation time 𝜏_𝑝_ of normal modes autocorrelation for modes 𝑝 = 1, 2,…, 𝑁 − 1 of E-K sequences as a function of chain length 𝑁 for different normalized sequence charge decoration nSCD values: (a) nSCD = 0, (b) nSCD = 0.02, (c) nSCD = 0.20, (d) nSCD = 0.46, and (e) nSCD = 1. Colors, ranging from cyan to magenta, represent increasing 𝑁.

**Figure S9:**
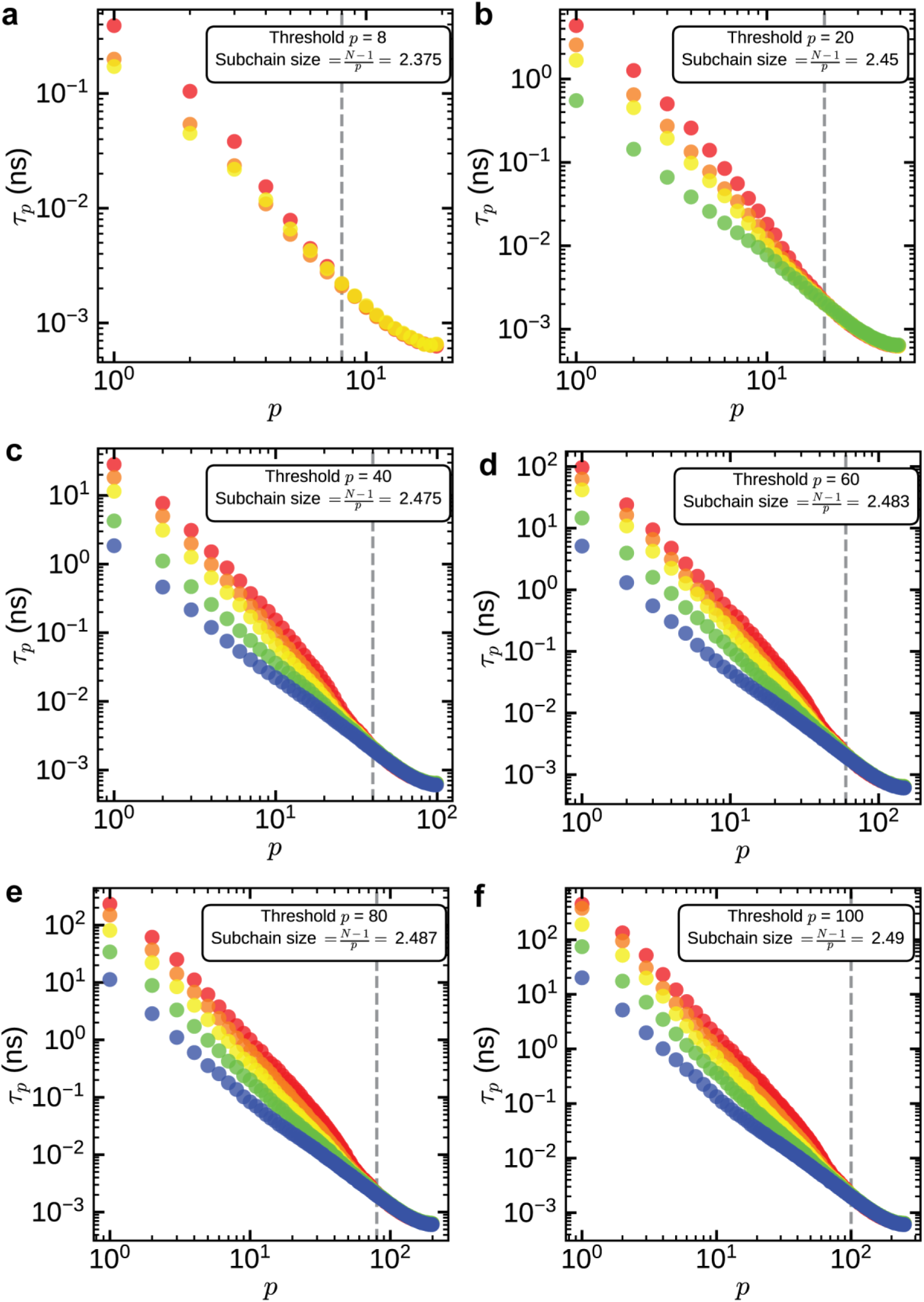
Relaxation Time of Normal Modes Autocorrelation for E-K Sequences **Within the Dense Phase:** Relaxation time 𝜏_𝑝_ of normal modes autocorrelation for modes 𝑝 = 1, 2,…, 𝑁 − 1 of E-K sequences as a function of normalized sequence charge decoration nSCD values for different chain lengths 𝑁: (a) 𝑁 = 20, (b) 𝑁 = 50, (c) 𝑁 = 100, (d) 𝑁 = 150, (e) 𝑁 = 200, and (f) 𝑁 = 250. Dashed lines represent the threshold mode number beyond which nSCD does not affect relaxation times. Symbol colors, ranging from blue to red, represent increasing nSCD values.

**Figure S10:**
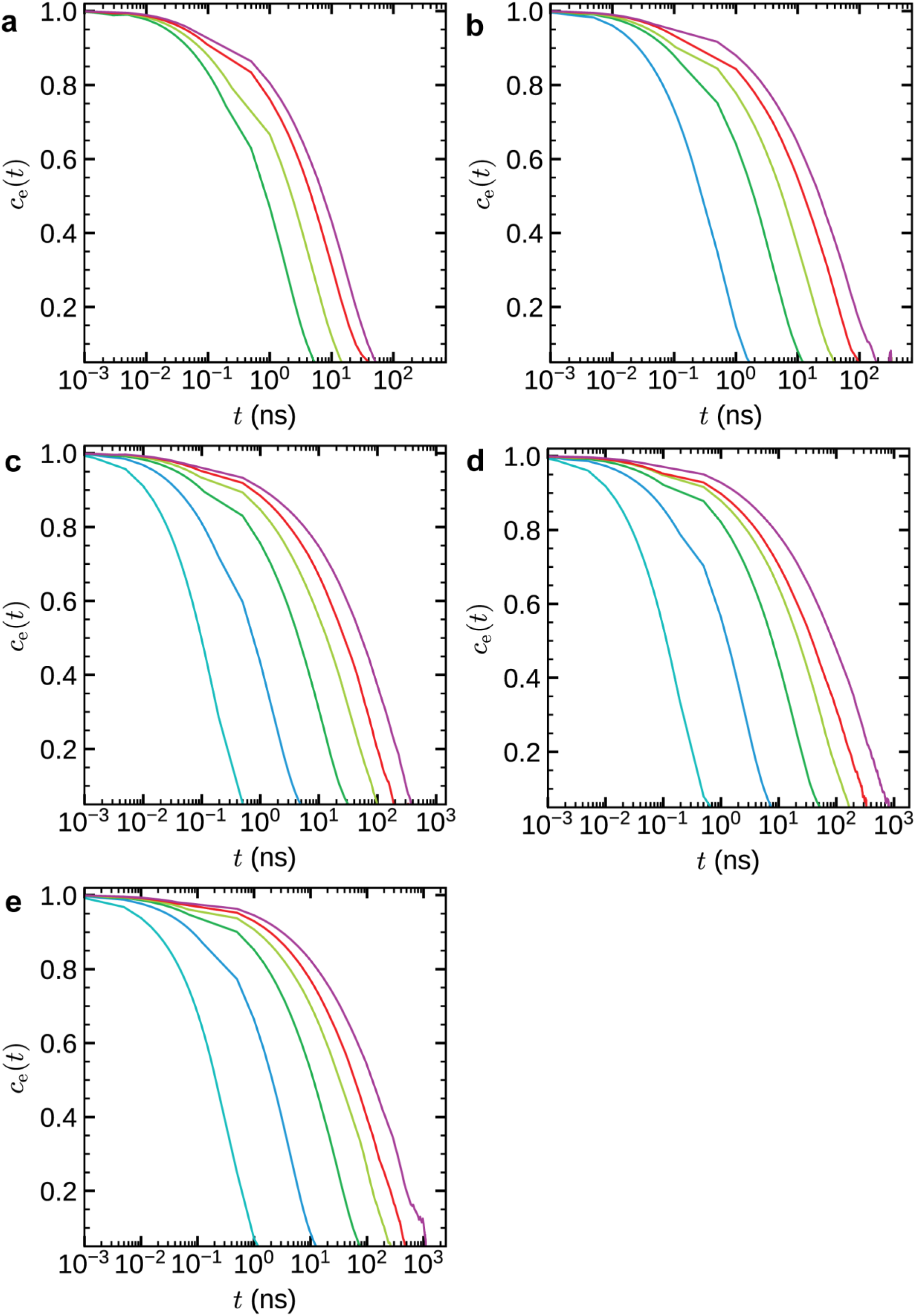
End-to-End Vector Autocorrelation Function for E-K Sequences Within the Dense Phase: End-to-end vector autocorrelation function 𝑐_e_(𝑡) of E-K sequences as a function of chain length 𝑁 for different normalized sequence charge decoration nSCD values: (a) nSCD = 0, (b) nSCD = 0.02, (c) nSCD = 0.20, (d) nSCD = 0.46, and (e) nSCD = 1. Colors, ranging from cyan to magenta, represent increasing 𝑁.

**Figure S11:**
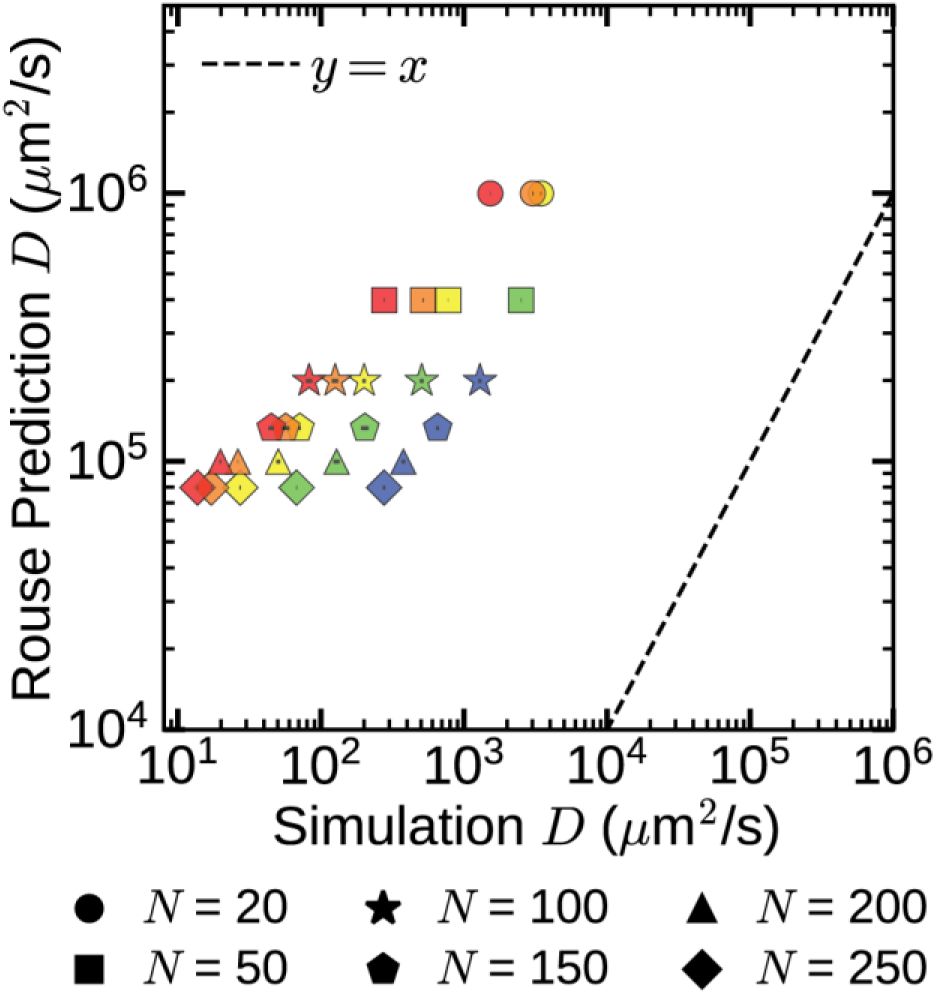
Rouse Theory Prediction of Dense Phase Chain Diffusivity of E-K Sequences Using Monomer Friction: Correlation plot comparing Rouse theory predictions and simulated dense phase chain diffusion coefficients for E-K sequences of varying chain length 𝑁 and normalized sequence decoration nSCD value, where predictions were made using Equation (2). The dashed line in the plot denotes the 𝑦 = 𝑥 line. Symbol colors, ranging from blue to red, correspond to increasing nSCD values.

**Figure S12:**
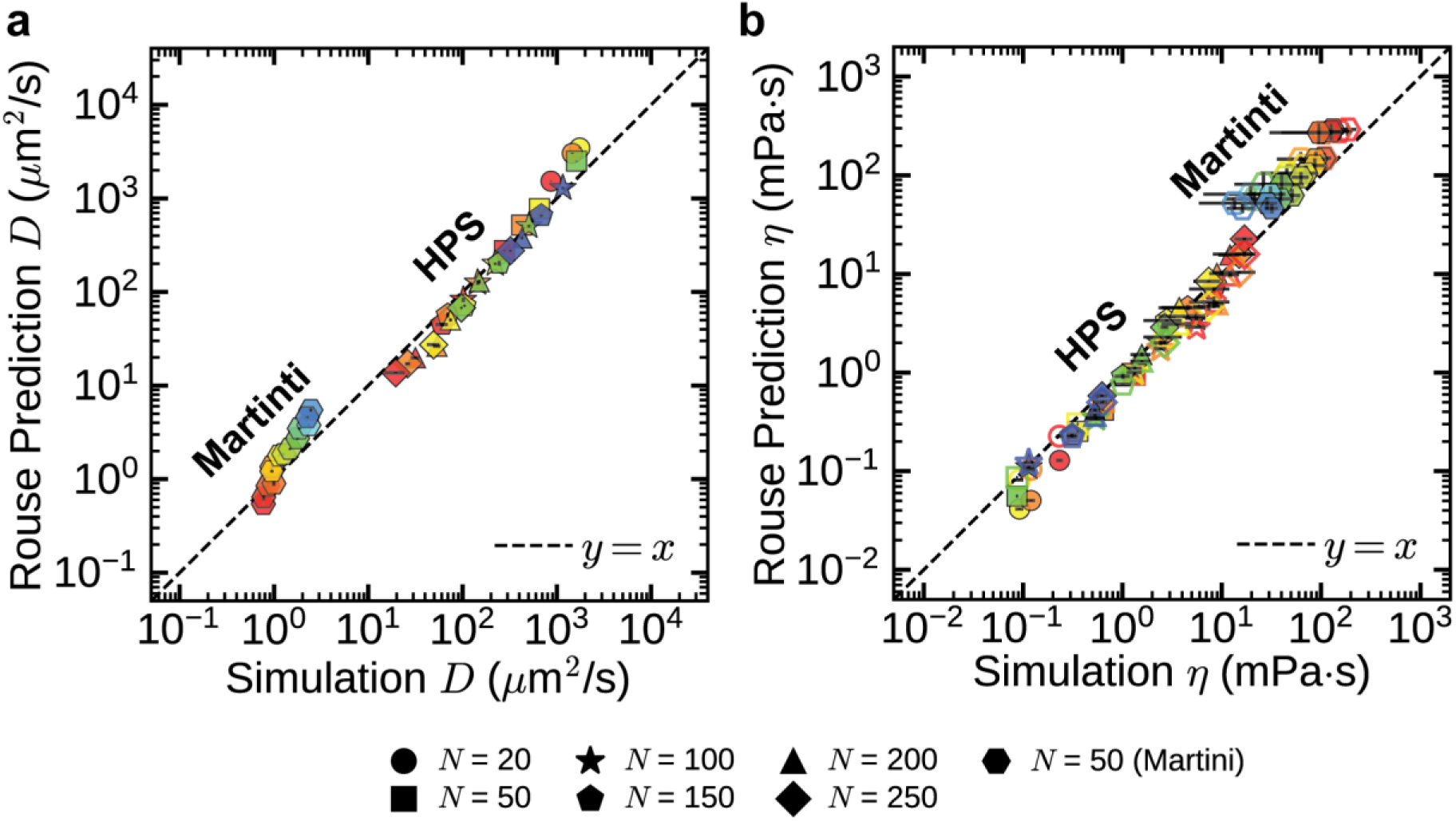
**Rouse Theory Prediction of Dense Phase Material Properties of E-K Sequences Using Chain Reconfiguration Times**: Correlation plots comparing Rouse theory predictions and simulated dense phase material properties for E-K sequences of varying chain length 𝑁 and normalized sequence charge decoration nSCD value, with (Martini) and without (HPS) explicit solvent interactions: (a) diffusivity 𝐷, predicted using equation (4), and (b) viscosity 𝜂. In (b), hollow markers indicate predictions using equation (5), and filled markers indicate those from equation (6). The dashed line in the plots denotes the 𝑦 = 𝑥 line. Symbol colors, ranging from blue to red, correspond to increasing nSCD values.

**Figure S13:**
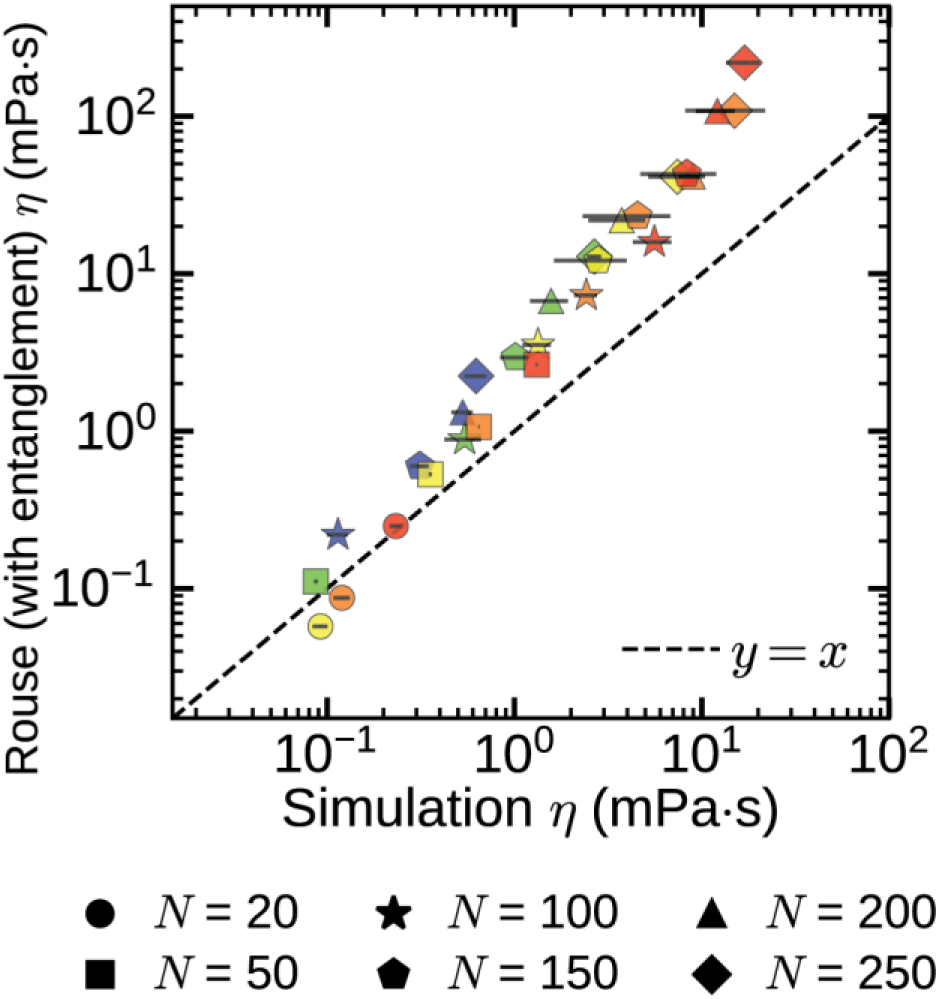
**Rouse Theory (With Entanglements) Prediction of Dense Phase Viscosity of E-K Sequences Using Chain Reptation Times**: Correlation plot comparing Rouse theory (with entanglements) predictions and simulated dense phase viscosities for E-K sequences of varying chain length 𝑁 and normalized sequence charge decoration nSCD value, where predictions were made using Equation (33). The dashed line in the plot denotes the 𝑦 = 𝑥 line. Symbol colors, ranging from blue to red, correspond to increasing nSCD values.

**Figure S14:**
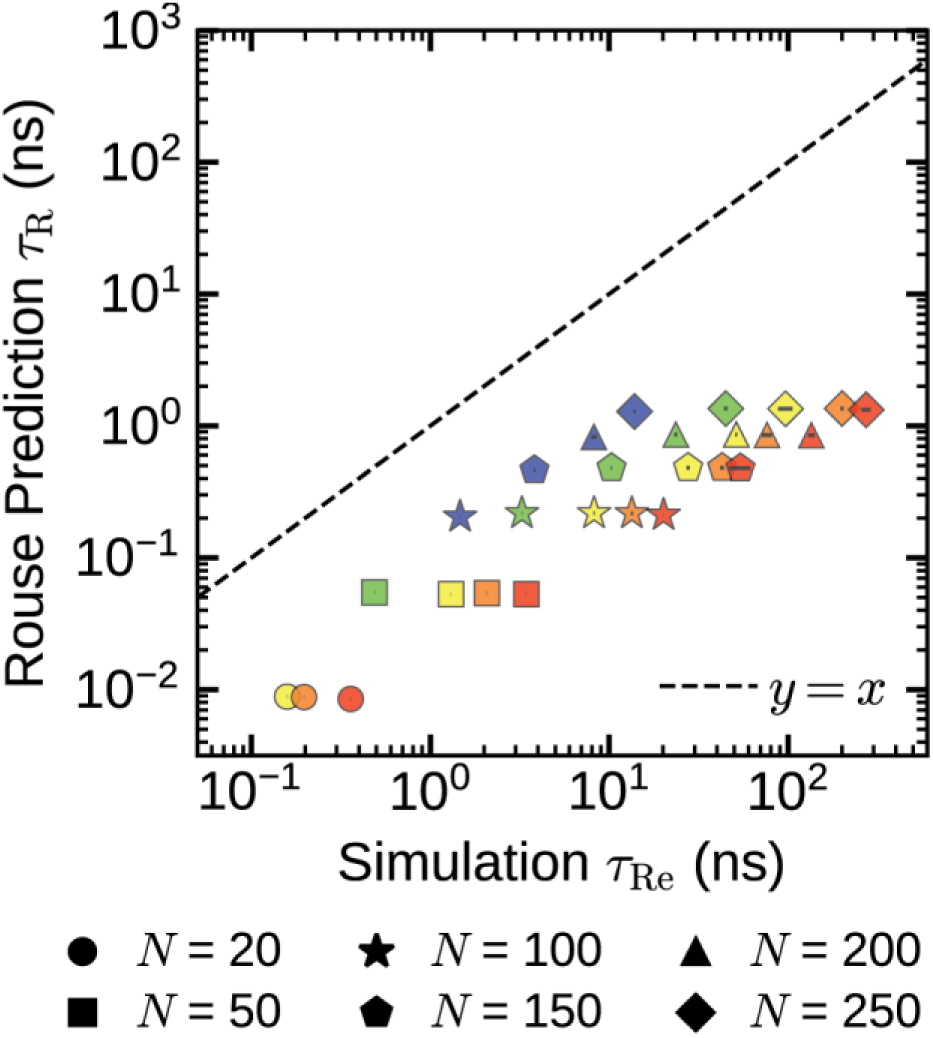
Rouse Theory Prediction of Dense Phase Chain Reconfiguration Times of E-K Sequences Using Monomer Relaxation Times: Correlation plot comparing Rouse theory predictions and simulated dense phase chain reconfiguration times for E-K sequences of varying chain length 𝑁 and normalized sequence decoration nSCD value, where predictions were made using Equation (7). The dashed line in the plot denotes the 𝑦 = 𝑥 line. Symbol colors, ranging from blue to red, correspond to increasing nSCD values.

**Figure S15:**
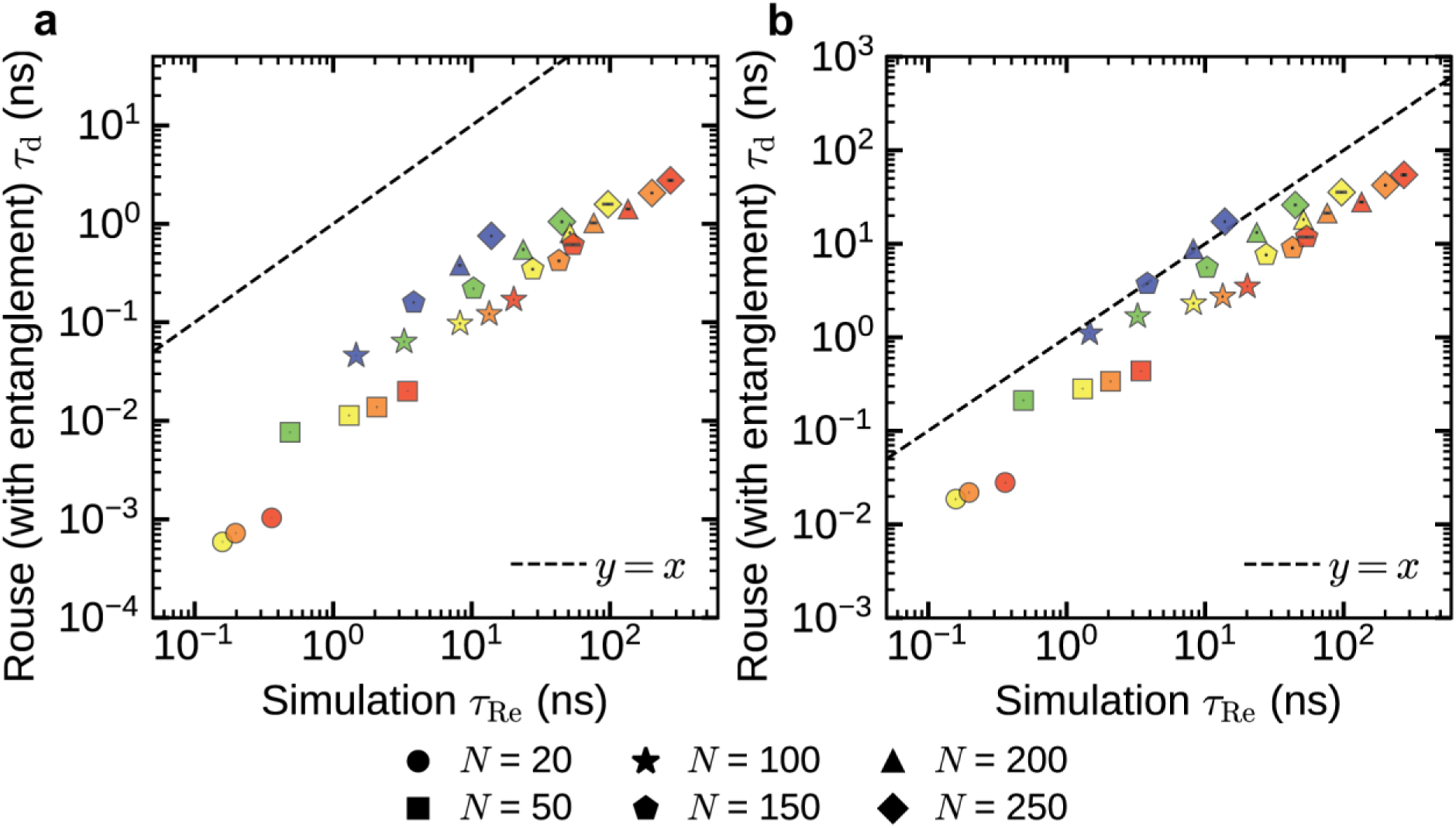
**Rouse Theory (With Entanglements) Prediction of Dense Phase Chain Reconfiguration Times of E-K Sequences**: Correlation plots comparing Rouse theory (with entanglements) predictions and simulated dense phase chain reconfiguration times for E-K sequences of varying chain length 𝑁 and normalized sequence charge decoration nSCD value where (a) predictions were made using Equation (30), and (b) predictions were made using Equation (31). The dashed line in the plots denotes the 𝑦 = 𝑥 line. Symbol colors, ranging from blue to red, correspond to increasing nSCD values.

**Figure S16:**
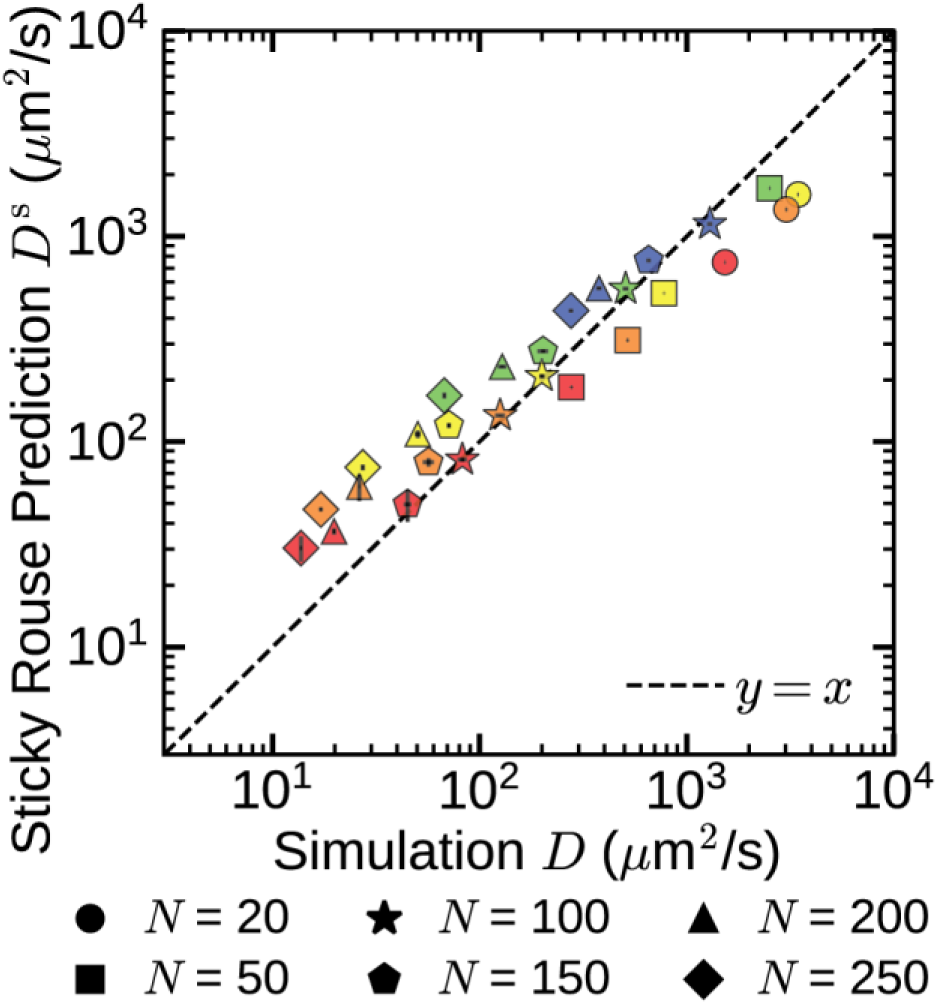
Sticky Rouse Theory Prediction of Dense Phase Chain Diffusivity of E-K Sequences Using Chain Reconfiguration Times: Correlation plot comparing sticky Rouse theory predictions and simulated dense phase chain diffusion coefficients for E-K sequences of varying chain length 𝑁 and normalized sequence decoration nSCD value, where predictions were made using predicted chain reconfiguration times 𝜏_R_^s^ in Equation (4). The dashed line in the plot denotes the 𝑦 = 𝑥 line. Symbol colors, ranging from blue to red, correspond to increasing nSCD values.

**Figure S17:**
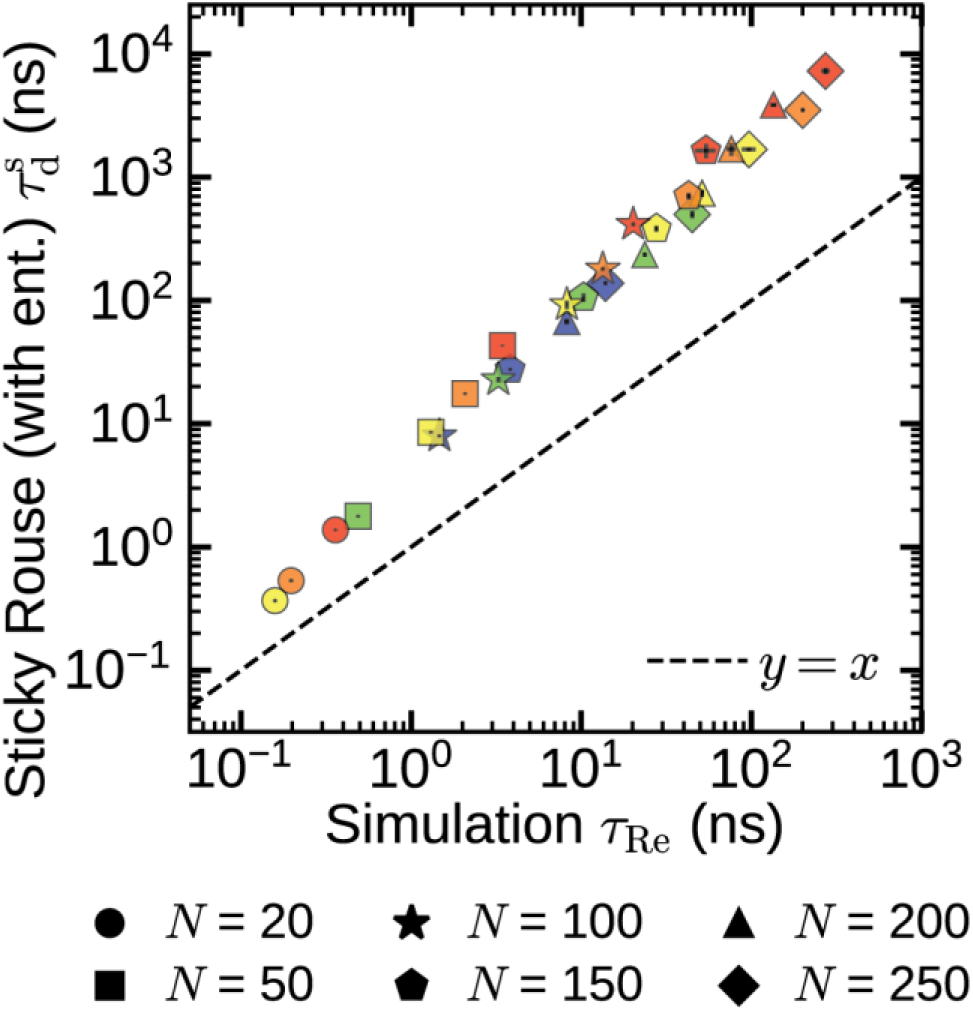
**Sticky Rouse Theory (With Entanglements) Prediction of Dense Phase Chain Reconfiguration Times of E-K Sequences**: Correlation plot comparing sticky Rouse theory (with entanglements) predictions and simulated dense phase chain reconfiguration times for E-K sequences of varying chain length 𝑁 and normalized sequence charge decoration nSCD value where predictions were made using Equation (35). The dashed line in the plots denotes the 𝑦 = 𝑥 line. Symbol colors, ranging from blue to red, correspond to increasing nSCD values.

**Figure S18:**
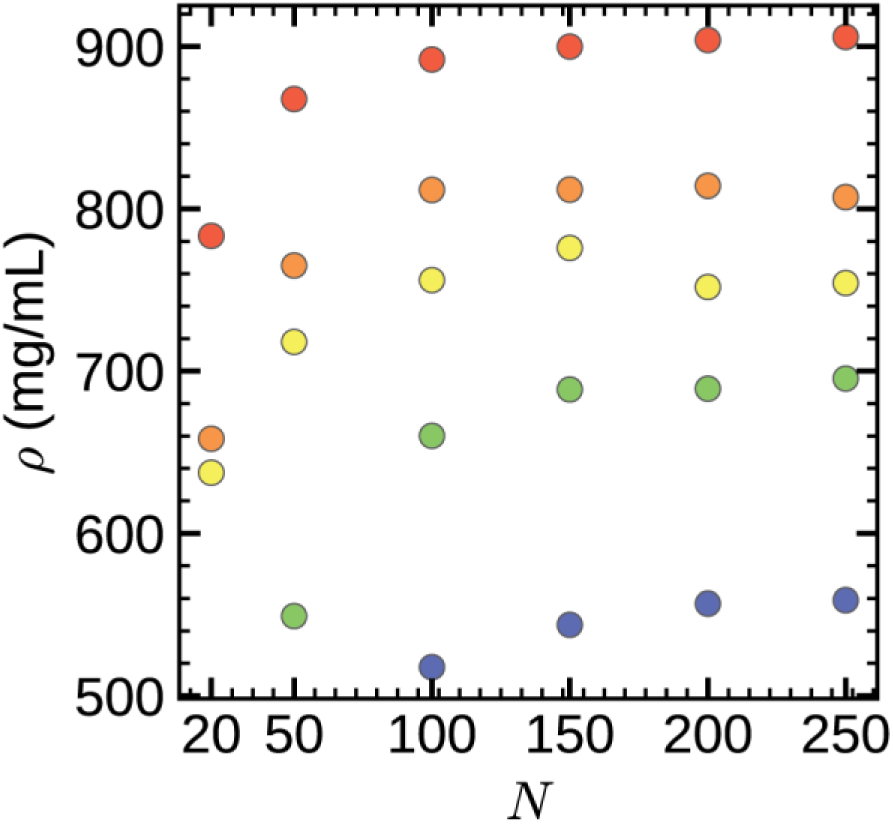
Dense Phase Concentration Attained by E-K Sequences in a Cubic Geometry During NPT Equilibration: Dense phase concentrations ρ attained by E-K sequences during the NPT equilibration as a function of chain length 𝑁 for different normalized sequence charge decoration nSCD values: **(a)** nSCD = 0, **(b)** nSCD = 0.02, **(c)** nSCD = 0.20, **(d)** nSCD = 0.46, and **(e)** nSCD = 1. Symbol colors, ranging from blue to red, indicate increasing nSCD values.

**Figure S19:**
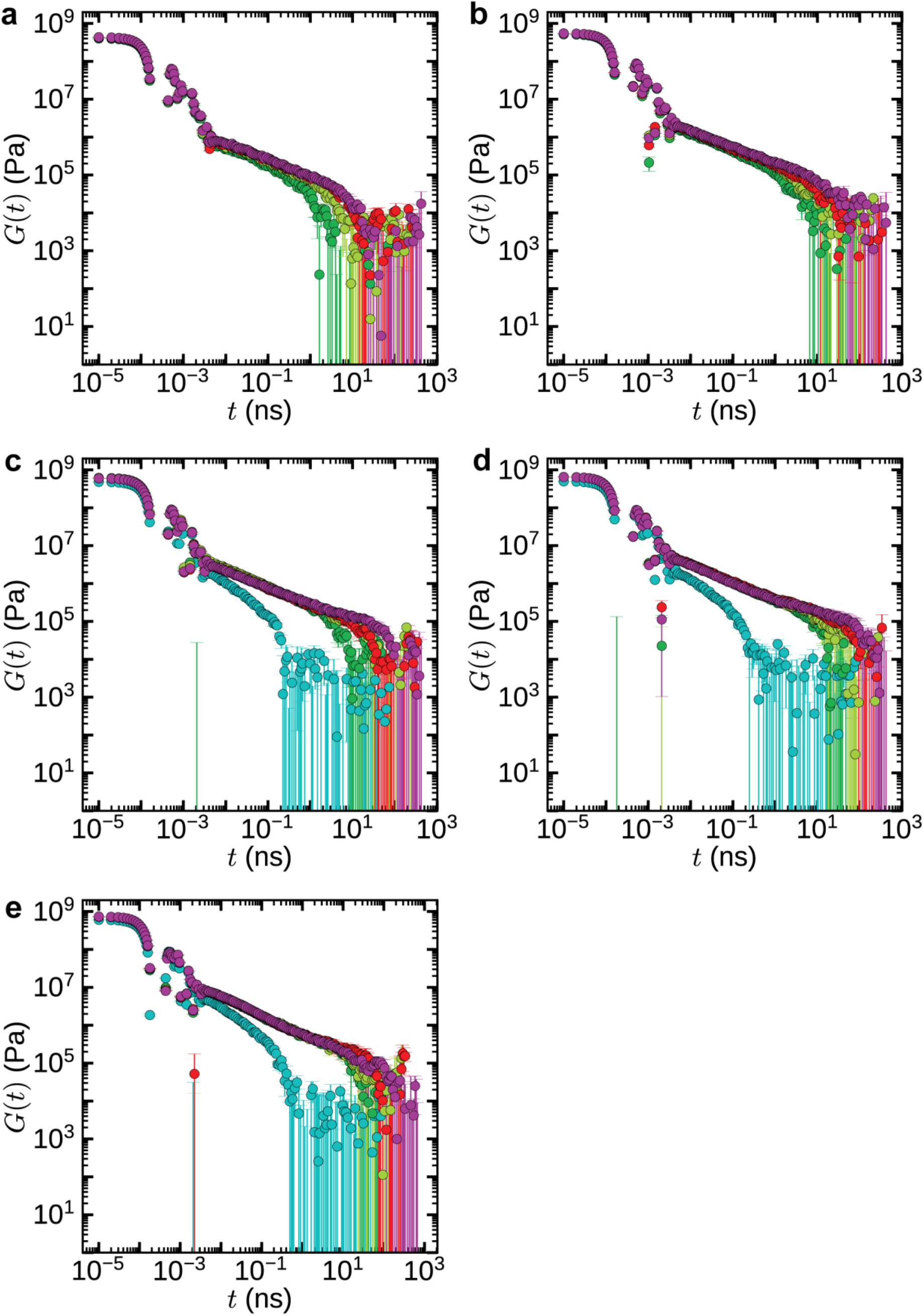
Shear Stress Relaxation Modulus of the Dense Phase Formed by E-K Sequences: Shear stress relaxation modulus 𝐺(𝑡) of the dense phase formed by E-K sequences as a function of chain length 𝑁 for different normalized sequence charge decoration nSCD values: (a) nSCD = 0, (b) nSCD = 0.02, (c) nSCD = 0.20, (d) nSCD = 0.46, and (e) nSCD = 1. Colors, ranging from cyan to magenta, represent increasing 𝑁.

**Figure S20:**
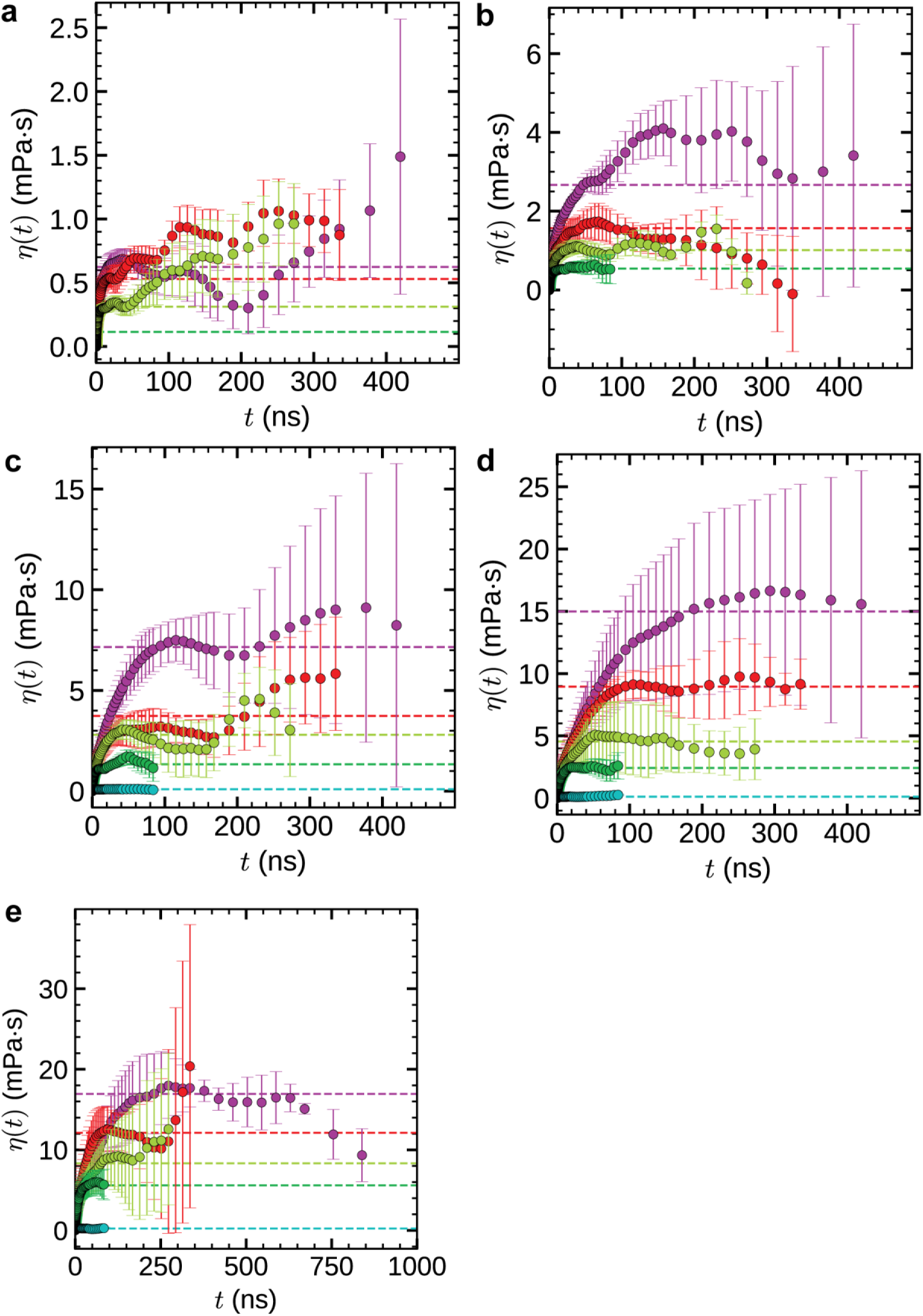
Bulk Viscosity of the Dense Phase Formed by E-K Sequences: Bulk viscosity 𝜂 of the dense phase formed by E-K sequences as a function of chain length 𝑁 for different normalized sequence charge decoration nSCD values: (a) nSCD = 0, (b) nSCD = 0.02, (c) nSCD = 0.20, (d) nSCD = 0.46, and (e) nSCD = 1. Horizontal dashed lines represent the average viscosity, calculated by averaging viscosity values in the plateau region. Colors, ranging from cyan to magenta, represent increasing 𝑁.

**Figure S21:**
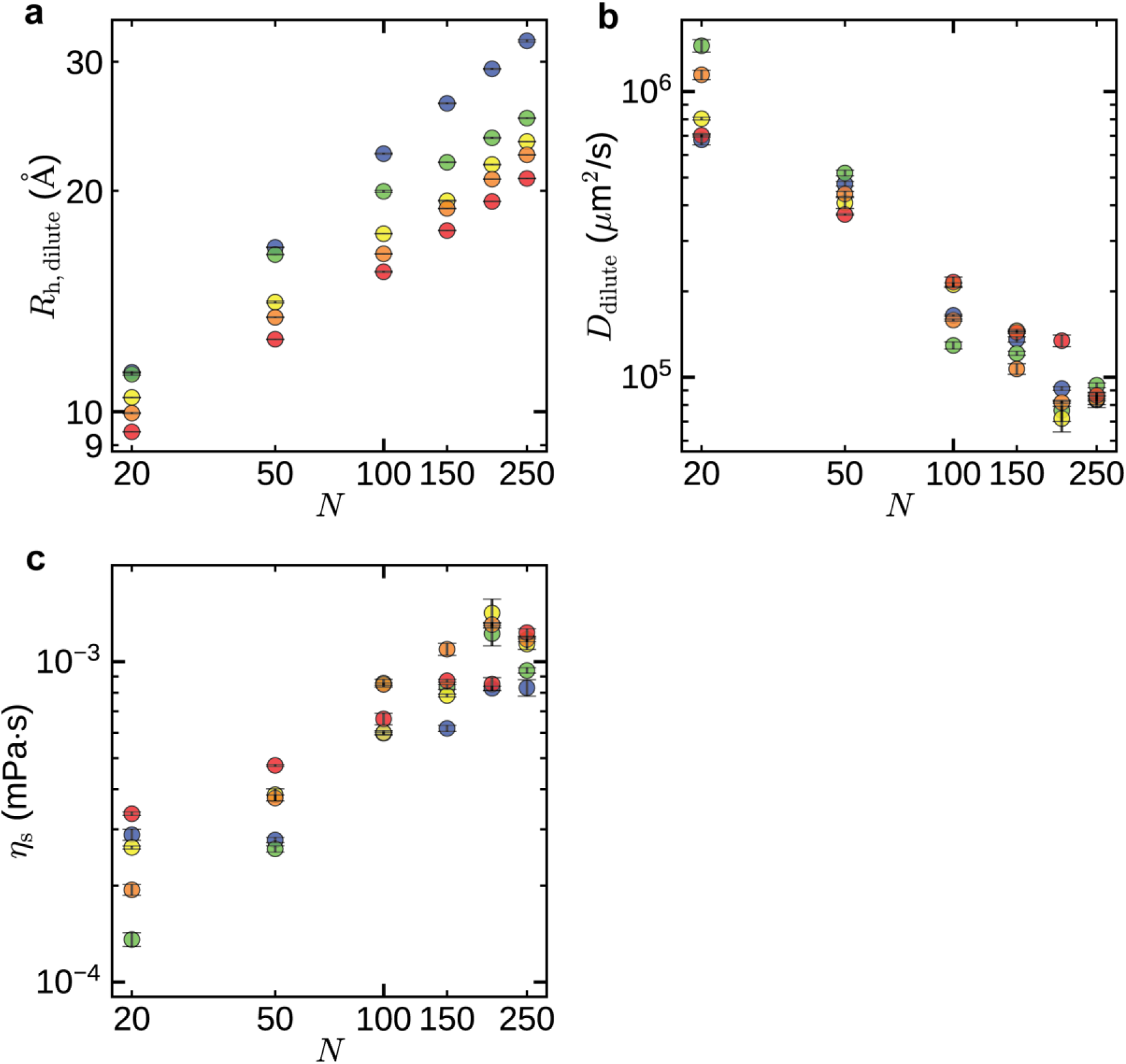
Hydrodynamic Radius, Diffusion Coefficient, and Solvent Viscosity of E-K Sequences in the Dilute Phase: (a) Hydrodynamic radius 𝑅_h,dilute_, (b) diffusion coefficient 𝐷_dilute_, and (c) solvent viscosity 𝜂_s_ in the dilute phase as a function of chain length 𝑁 and normalized sequence charge decoration nSCD value. Symbol colors, ranging from blue to red, represent increasing nSCD values.

**Figure S22:**
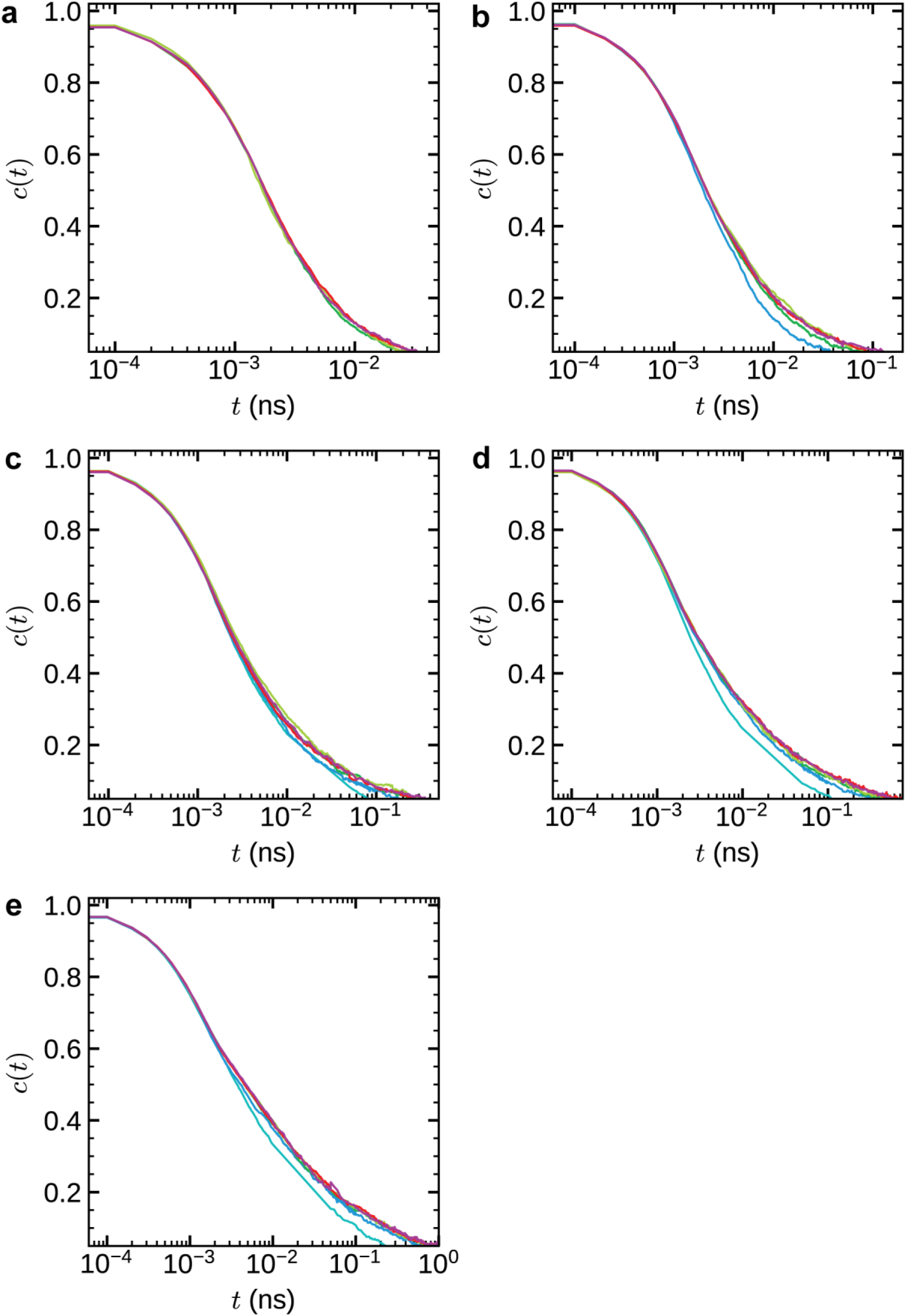
Intermittent Contact Lifetime Autocorrelation Function for E-K Sequences Within the Dense Phase: Intermittent contact lifetime autocorrelation function 𝑐(𝑡) of E-K sequences as a function of chain length 𝑁 for different normalized sequence charge decoration nSCD values: (a) nSCD = 0, (b) nSCD = 0.02, (c) nSCD = 0.20, (d) nSCD = 0.46, and (e) nSCD = 1. Colors, ranging from cyan to magenta, represent increasing 𝑁.

**Figure S23:**
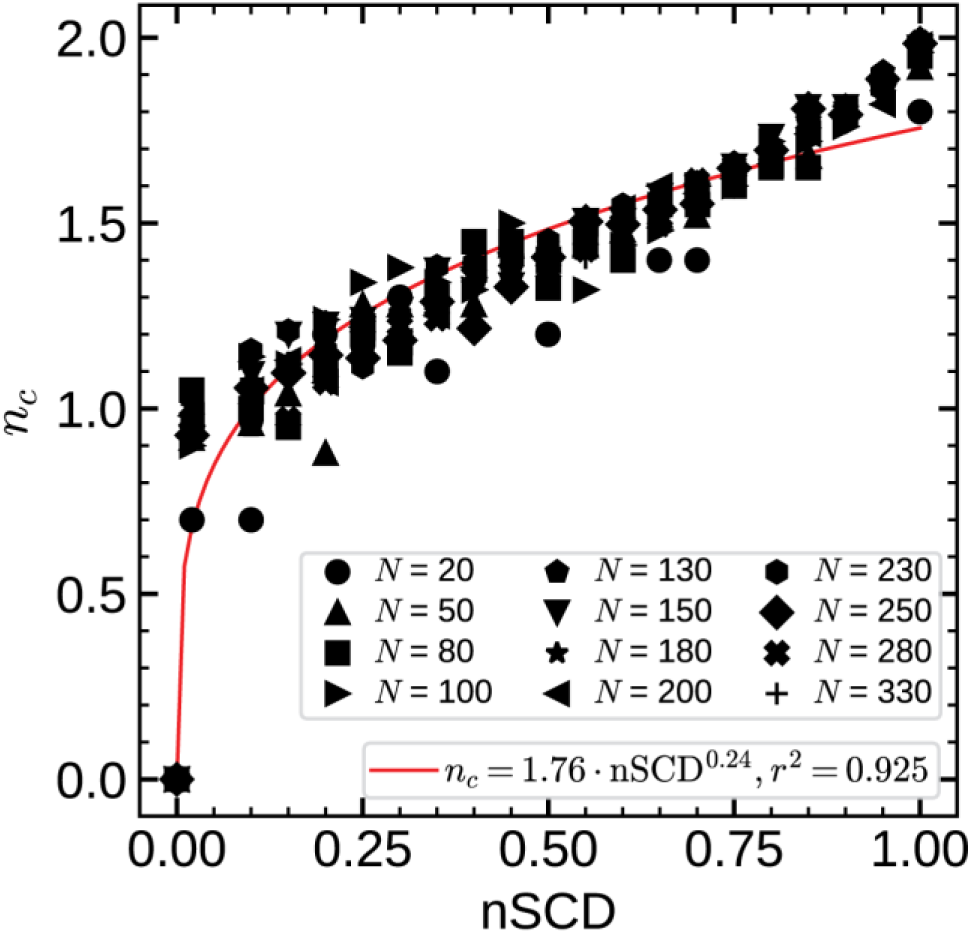
Average Number of Immediate Neighbors with the Same Charge in E-K Sequences: Average number of immediate neighbors (left and right) with the same charge, 𝑛_𝑐_, across the sequence in E–K sequences, shown as a function of chain length *N* and normalized sequence charge decoration nSCD values. The solid line represents the power law fit.

**Figure S24:**
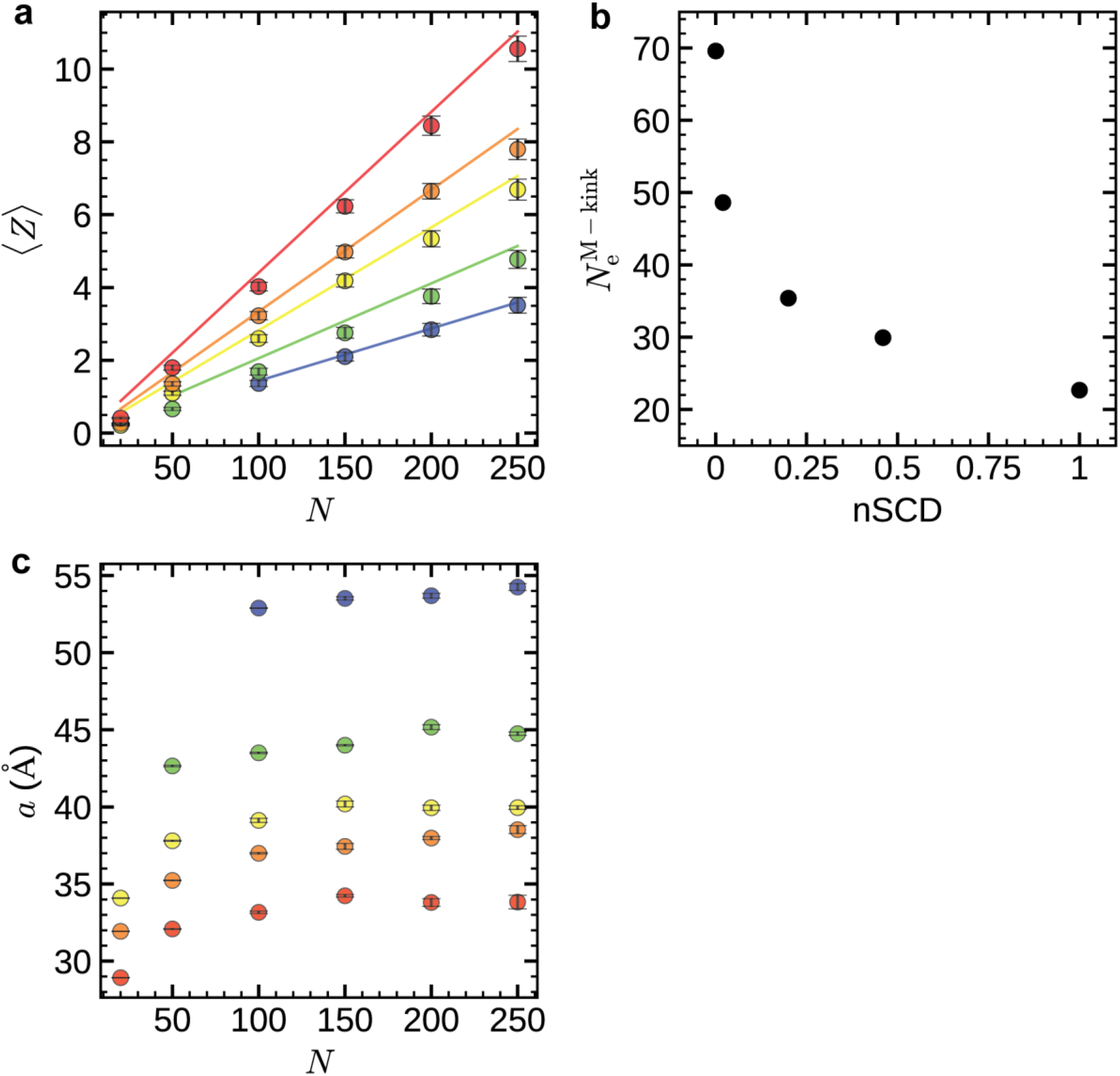
Calculation of Entanglement Spacing Within the Dense Phase for E-K Sequences: (a) Average number of entanglements (kinks) per chain ⟨𝑍⟩, (b) entanglement length 𝑁_e_^M−kink^, and (c) entanglement spacing 𝑎 of E-K sequences within the dense phase as a function of chain length 𝑁 and normalized sequence charge decoration nSCD value. The solid lines in (a) represent the ⟨𝑍⟩ = 𝑁^M-kink^𝑁 fits. Symbol colors in (a) and (c), ranging from blue to red, represent increasing nSCD values.

**Figure S25:**
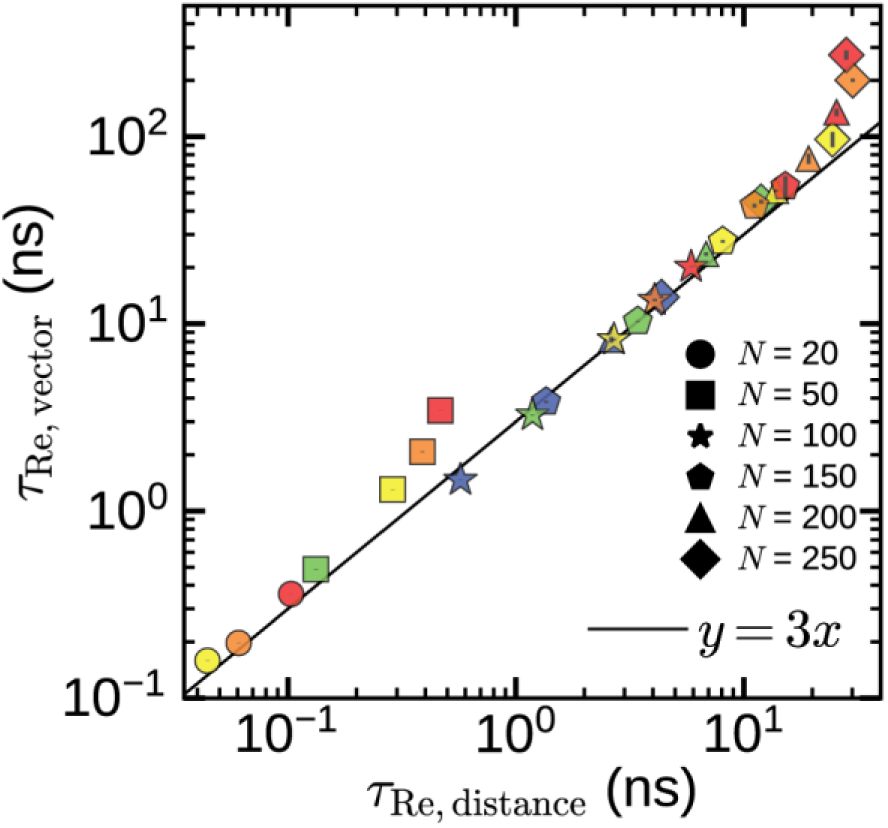
Comparison of End-to-End Vector and Distance Relaxation Times of E-K sequences Within the Dense Phase: Correlation plot comparing the end-to-end vector relaxation times 𝜏_Re, vector_ and distance relaxation times 𝜏_Re, distance_ of E-K sequences (computed from simulations) within the dense phase as a function of chain length 𝑁 and normalized sequence charge decoration nSCD value. The solid line represents 𝑦 = 3𝑥 reference line. Symbol colors, ranging from blue to red, represent increasing nSCD values.

---

**Table 1:**
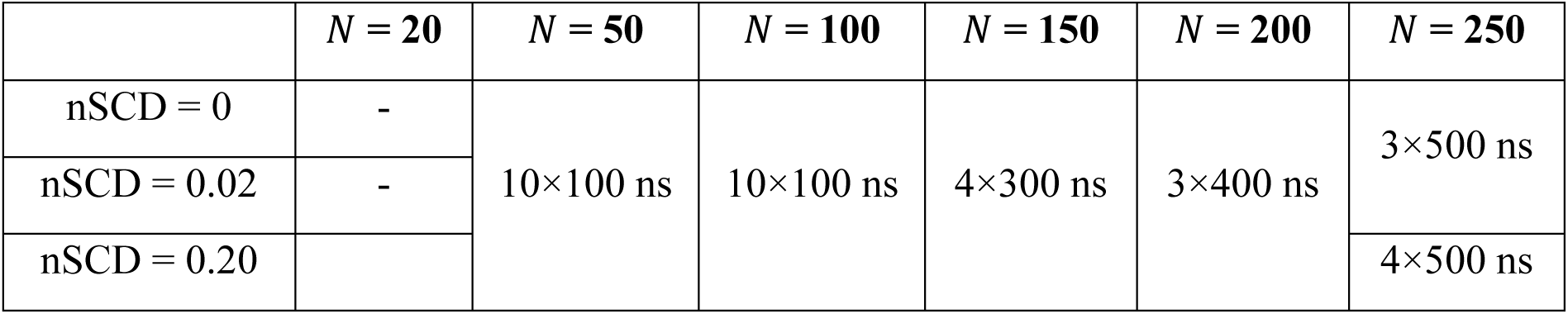
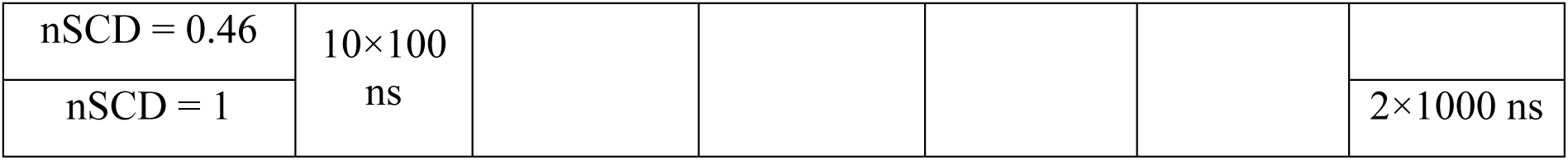
Simulation details for dense phase viscosity calculations: Simulation durations for the NVT runs performed to compute the zero-shear viscosity 𝜂 of E-K sequences for different chain lengths 𝑁 and normalized sequence charge nSCD. Each entry indicates the number of independent simulations 𝑛 and their duration 𝑡 (i.e., 𝑛 × 𝑡).

**Supplementary Table S1:**
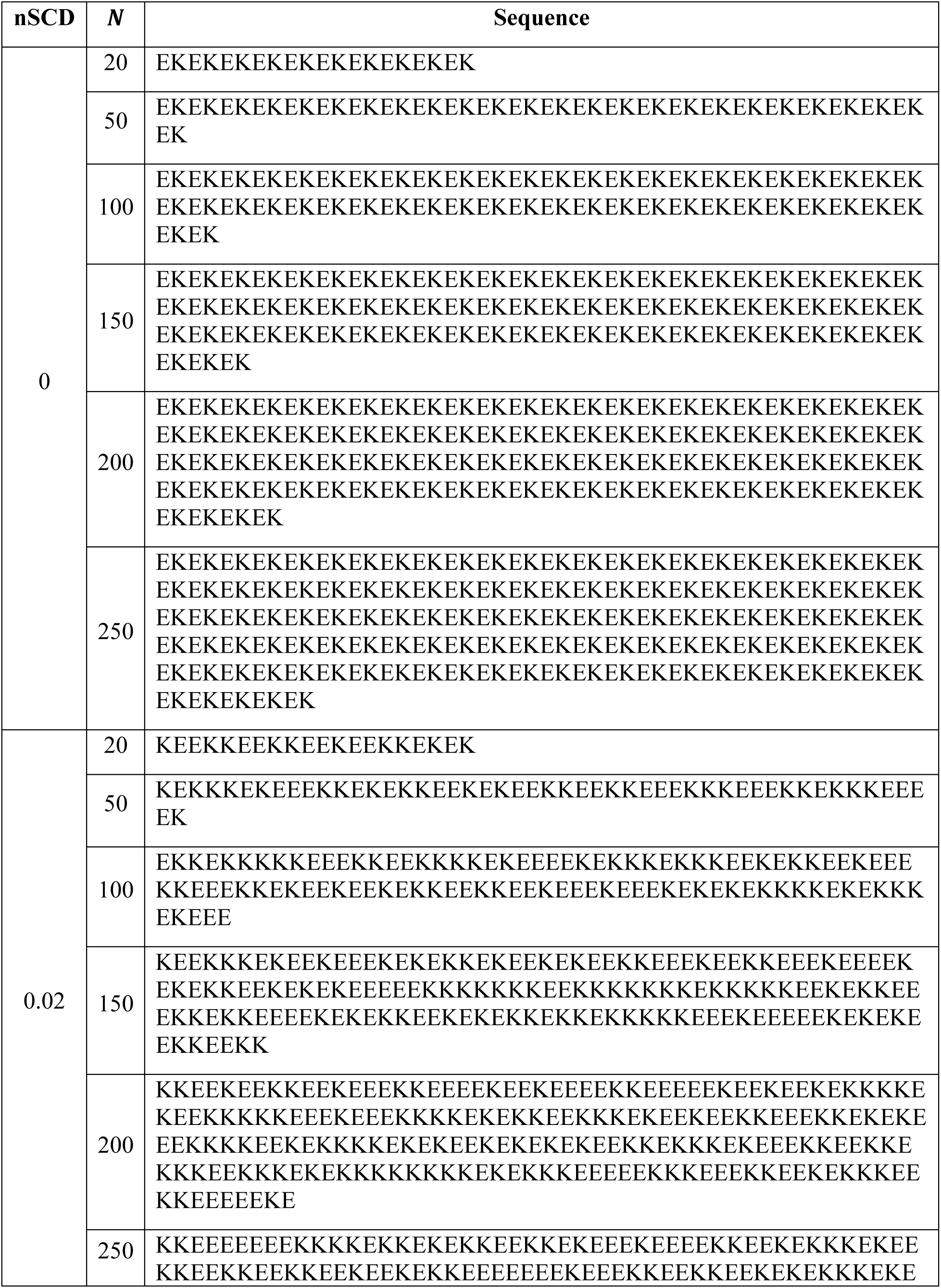
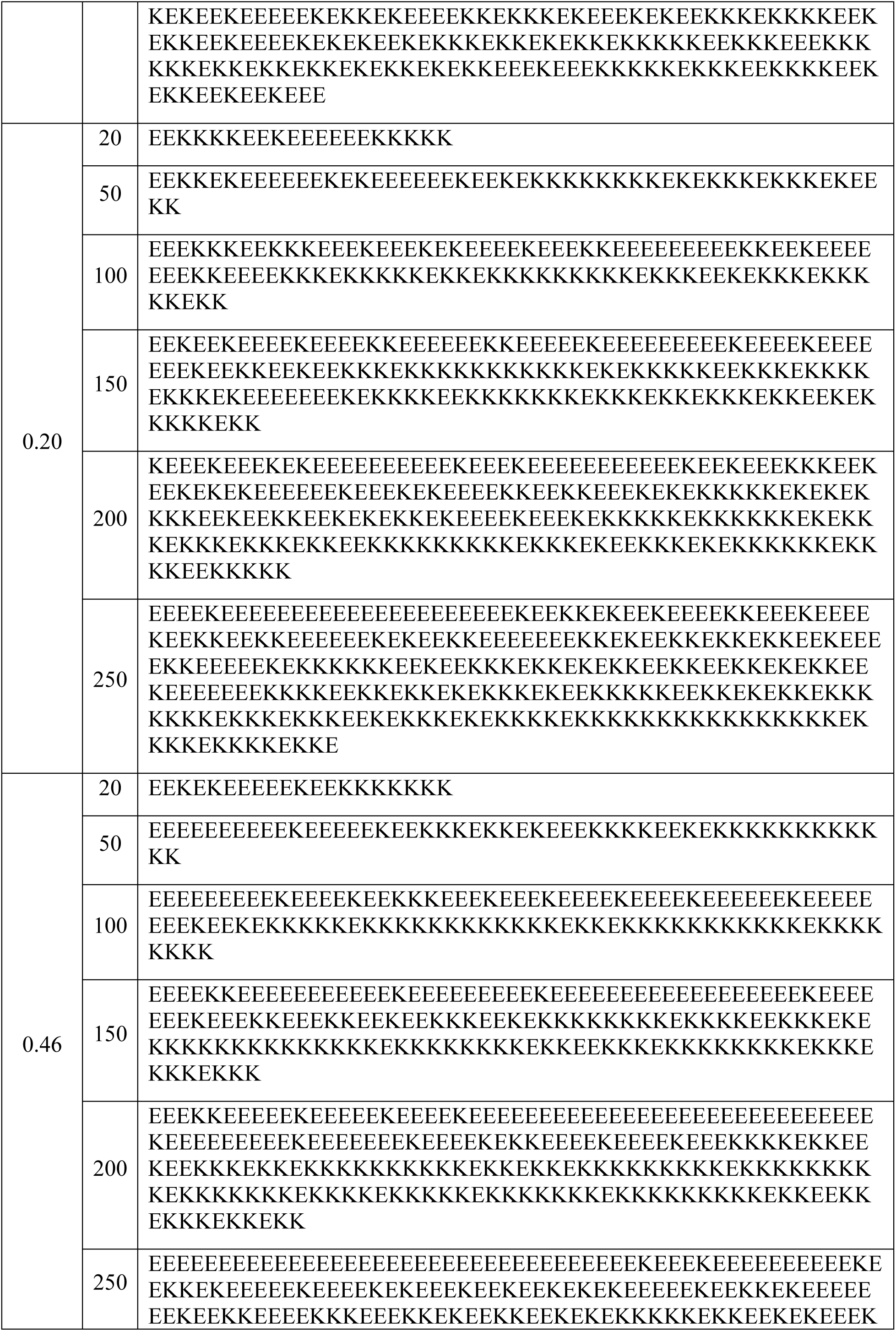
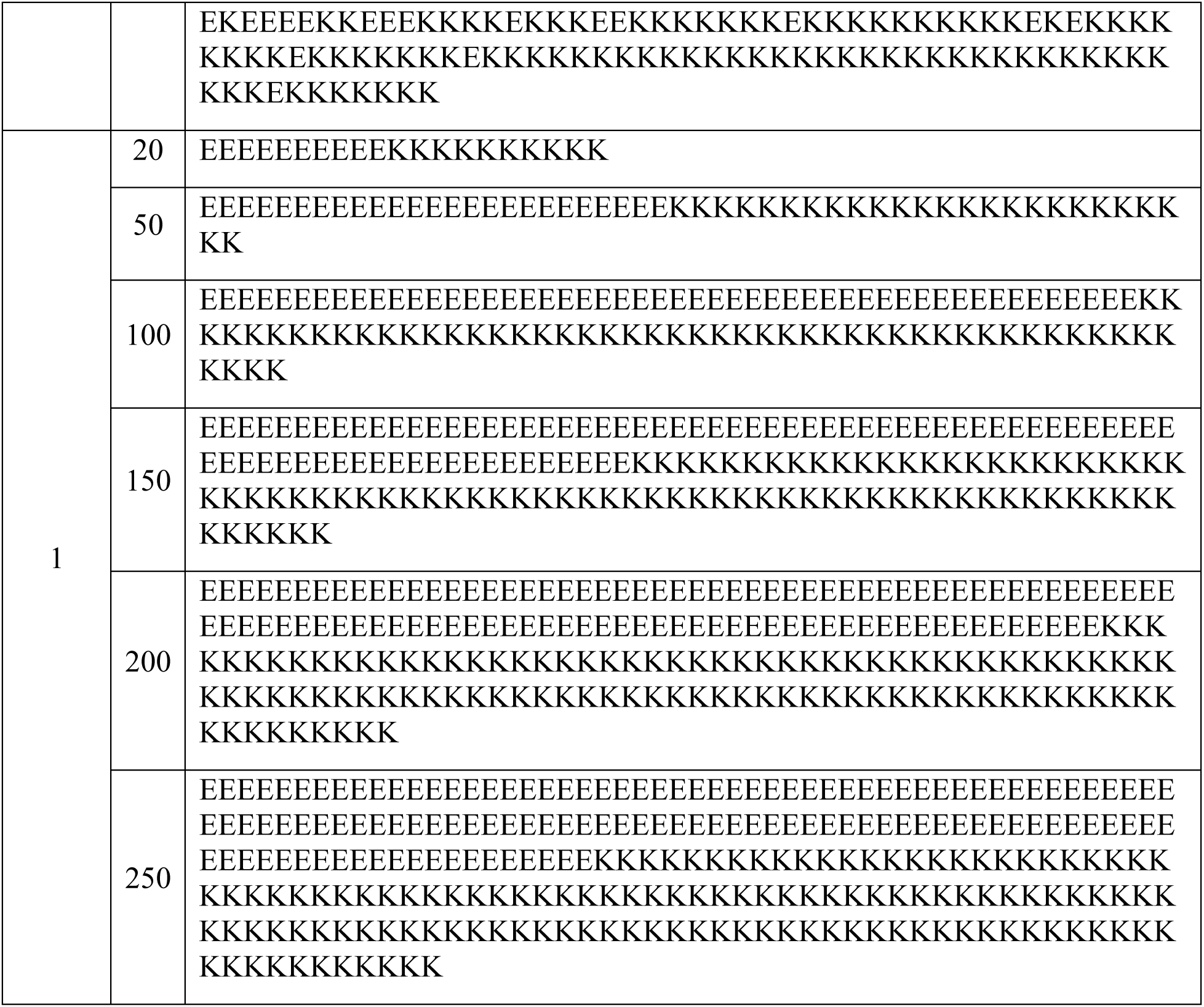
Amino Acid Arrangements for E-K Sequences Used in This Study: Table listing the specific amino acid arrangements of E-K sequences utilized in the simulations, categorized by chain length 𝑁 and charge patterning nSCD values.

